# Unraveling HIV protease drug resistance and genetic diversity with kernel methods

**DOI:** 10.1101/2025.03.26.644092

**Authors:** Elies Ramon

## Abstract

A definitive cure for HIV/AIDS does not exist yet and, thus, patients rely in antiretroviral therapy for life. In this scenario, the emergence of drug resistance is an important concern. The automatic prediction of resistance from HIV sequences is a fast tool for physicians to choose the best possible medical treatment. This paper proposes three kernel functions to deal with this data: one focused on single residue mutations, another on *k*-mers (close-range information in sequence), and another on pairwise interactions between amino acids (close and long-range information). Furthermore, the three kernels are able to deal with the categorical nature of HIV data and the presence of allelic mixtures. The experiments on the PI dataset from the Stanford Genotype-Phenotype database show that they generate prediction models with a very good performance, while remaining simple, open and interpretable. Most of the mutations and patterns they consider relevant are in agreement with previous literature. Also, this paper compares the different but complementary view that two kernel methods (SVM and kernel PCA) give over HIV data, showing that the former is focused on optimizing prediction while the latter summarizes the main patterns of genetic diversity, which in the Stanford Genotype-Phenotype database are related to drug resistance and HIV subtype.

## 1 Introduction

Acquired immunodeficiency syndrome (AIDS) is a deathly condition caused by the human immunodeficiency virus (HIV). This retrovirus has a high prevalence worldwide: in 2023, 39.9 million people globally had HIV. The same year, 1.3 million people became newly infected, and 630,000 people died of AIDS-related illnesses [UNAIDS, 2024]. A definitive cure for the infection does not exist yet and, thus, patients rely on antiretroviral therapy for life. This treatment is based on a cocktail of two or three specific drugs that target key enzymes for the virus replication cycle: protease inhibitors (PIs), reverse transcriptase inhibitors (NRTIs and NNRTIs), integrase inhibitors (INIs) and fusion inhibitors. Current therapy has been so successful that, in many cases, viral load can be effectively suppressed and the patient never progresses to AIDS. However, resistance occurs in all antiretrovirals and can pose a threat to treatment effectiveness. The emergence of resistant variants is caused by HIV’s high mutation rate (10^*−*4^ mutations per nucleotide site per replication cycle), high recombination rate (10^*−*5^ − 10^*−*4^ breakpoints per site per generation) [Song et al., 2018], short replication time (1-2 days) and high virion yields (10^8^ − 10^9^ per day) [Iyidogan and Anderson, 2014]. Those traits allow a fast exploration of potential genetic variants to overcome selective pressure, like host immune system or drug therapy. When left unchecked, HIV evolves very fast within their hosts and creates complex and dynamic constellations of distinct but genetically linked variants: the quasispecies [Iyidogan and Anderson, 2014, Wensing et al., 2010]. Resistance can occur if the virus replication is not effectively suppressed (for instance, due to incomplete adherence to treatment) and these resistant variants can be transmitted to other hosts. Not all drugs present the same genetic barrier to resistance. In some drugs, a single mutation may be enough, while in others, HIV needs accumulating multiple mutations before becoming resistant [Duffey et al., 2024]. Some may affect a single inhibitor within a drug class, while others confer cross-resistance. The accumulation of mutations leads to complex patterns that become strongly entrenched; this explains why drug-resistant HIV variants are unlikely to revert [Zhang et al., 2020].

In this scenario, genotypic prediction from HIV sequences is a fast tool for physicians to choose the best possible medical treatment. This strategy, which is especially useful in case of treatment failure, reduces patients’ exposure to suboptimal drug regimens. Furthermore, it is more widely available than the slower and more expensive *in vitro* phenotype resistance tests, which need specialized and centralized wet lab facilities [Duffey et al., 2024, Parkin et al., 2023]. The first genotypic prediction models that allowed to infer resistance from sequence data were rule-based. They stemmed from the observation that mutations of specific residues greatly increased the resistance to certain drugs. These are known as “major” or “primary” mutations. Instead, “minor” or “accessory” mutations confer some incremental resistance and/or restore the enzyme activity and viral fitness disrupted by major mutations. Finally, there exist neutral mutations not related to drug resistance [Haq et al., 2009]. Indeed, it has been observed that, when acquiring resistance to a drug, major mutations tend to be selected first and minor mutations tend to follow them; however, the difference between the two is not always clear-cut [Wensing et al., 2022]. In any case, the clarity and interpretability of rule-based models allowed studying the functional and structural changes induced by some well-known mutations. Furthermore, some mutational patterns (involving two or more drug-associated mutations) have also been studied, acknowledging potential interactions on top of the “additive” effects of point mutations. Rule-based estimates of resistance continue to be used to this day (see for instance the Stanford’s HIVdb system presented by Liu and Shafer [2006]). However, HIV evolves very rapidly, making it unfeasible to catalog all the new resistance mutations and mutational patterns that continually emerge.

It is precisely this enormous genetic variability what makes machine learning methods (ML), *e*.*g*. Linear Regression, Artificial Neural Networks, Random Forests and Support Vectors Machines (SVMs), so useful for the problem of HIV drug resistance prediction. The most simple ML approaches are sequence-based, *i*.*e*. each position (or mutation) is considered a feature. No prior association between sequence residues is assumed explicitly, so the assumption is that all residues mutate independently of one another. More ambitious studies that complement this information with structural and association data also exist [Bonet, 2015]. ML models have achieved very good performances in HIV resistance prediction, but they have their own set of weaknesses. In some cases, the interpretability that characterizes rule-based algorithms is lost. Also, the very nature of HIV sequences poses some challenges to training ML models. First of all, sequence data can be considered as a vector of categorical features, instead the classic real-valued vectors that are the preferred input for most ML methods. To bridge this gap, usually some kind of data pre-processing is performed. A second challenge is amino acid mixtures. Mixtures in this data indicate the presence of quasispecies [Iyidogan and Anderson, 2014], *i*.*e*. two or more virus variants within a patient. This poses difficulties for most ML and rule-based methods and, at the same time, obscures the genotype-phenotype correlation [Schmidt et al., 2000]. Typical pre-processing approaches entail keeping only one amino acid of the mixture or excluding the affected sequences, especially if the mixture is at a major drug resistance position. In fact, the latter stance is the one recommended by Stanford HIVdb. However, in some datasets, especially when dealing with therapy-experienced patients, HIV mixtures are so pervasive that a large fraction of sequences are affected.

In a previous work, we presented a ML approach that acknowledges amino acid mixtures as well as the categorical aspect of the HIV sequence data [Ramon et al., 2019]. We achieved a very good performance with our approach, which was based on SVMs coupled with nonlinear kernel functions specific for this data. Nevertheless, there were some blind spots our work did not address. For instance, we did not investigate which residues and/or mutations the SVM models considered important. On the contrary: we weighted each position with “a priori” importances computed by a Random Forest model. Although it is clear that some positions are key in acquiring resistance, not all amino acid substitutions are equally important, as acknowledged by catalogs of resistance-associated mutations. Moreover, weighting was not of much use in the PI models (although it dramatically improved prediction in most INIs and reverse transcriptase inhibitors models). At the end, by not making interpretable SVM models, we left out valuable information regarding why these models work well, and if they can be used for achieving relevant biological conclusions. Another limitation of our work was that we performed a sequence-based prediction without taking into account potential close and/or long range interactions between protein residues. In the case of HIV, it is known that the high rate of recombination greatly accelerates the disappearance of genetic linkage. However, other non-random associations between alleles or mutations may arise because of structural and functional interactions among protein residues. In this regard, Wang and Lee [2007] found that most associations detected in HIV data from therapy-experienced patients vanish in sequences from untreated HIV carriers, and that pairwise associations within the pol gene (which encodes protease, reverse transcriptase and integrase) are mainly due to functional interactions when the virus is under selective pressure. Unlike genetic linkage, functional associations can be found among distant positions at the sequence level (though, once the protein has folded, these residues may be spatially close). It is widely known that three-dimensional structure is key for enzymatic activity, as it enables substrate specificity and determines which reactions the enzyme will catalyze. Also, structure is often more conserved than protein sequence Rajendran et al. [2023]. Understanding protein structure and dynamics has been (and continues to be) of the utmost importance for rational drug design. PIs are a textbook example of this phenomenon: they are competitive inhibitors that mimic the transition state of HIV protease natural substrates, and were considered the first success of structure-based drug design [De Clercq, 2009].

HIV protease is enzymatically active as a homodimer Rajendran et al.-2023]. A big part of the dimer interface is stabilized by intermolecular contacts between N- and C-terminal regions (residues 1–5 and 95–99). During dimerization, both subunits interact at the active site interface. The process in completed in a second step, when further interactions between the terminal regions generate a stable protease [Hayashi et al., 2014]. The substrate binding pocket spans (approximately) residues 25-33, 47-53 and 80-84 [Iyidogan and Anderson, 2014, Ali et al., 2010] and is mostly hydrophobic in nature. Within the active site there is the conserved catalytic triad D25, T26 and G27. Also, a flexible glycine-rich flap region (residues 43-58) regulates the access of ligands (substrate and/or inhibitor) to the active site pocket. The flap hinge and elbow (35-42), fulcrum (10–23), and cantilever regions (59–75) are important for the opening of the flaps and flap dynamics [Dakshinamoorthy et al., 2023, Rajendran et al., 2023]. Ligand binding in protease involves conformational changes in the protease flaps and the hydrophobic core (which also includes residues like 76, 77, 85, 89, 90 and 93 [Seema et al., 2012]). During the resistance emergence, major mutations arise at the protease active site and its surroundings, expanding the pocket and reducing the binding free energy of the PIs. As these mutations often cause conformational changes, additional minor mutations are selected in turn to compensate the impaired catalytic activity and stabilize the enzyme structure [Iyidogan and Anderson, 2014, Dakshinamoorthy et al., 2023]. Some authors have defined 17 major drug resistance positions in protease (residues 23, 24, 30, 32, 33, 46, 47, 48, 50, 53, 54, 73, 76, 82, 84, 88, 90) and 24 minor positions (residues 10, 11, 13, 20, 34, 35, 36, 43, 45, 55, 58, 60, 63, 71, 74, 75, 77, 79, 83, 85, 89, 91, 93, 95) [Haq et al., 2009]. As the wild type (WT) protease is 99 residues long, this entails that at least a 40% of the sequence has some role in drug resistance, more than any other HIV target enzyme [Ali et al., 2010]. Also, unlike reverse transcriptase or integrase, resistance mutations are distributed along the whole sequence [Iyidogan and Anderson, 2014]. In real-life datasets, the most conserved domains are the N- and C-terminal regions, as well as the catalytic triad.

The main goal of this paper is using specific kernel functions for HIV data to unravel key mutations and patterns that enable drug resistance. More specifically, the points studied will be: (i) the relationship between HIV genetic diversity and resistance in the Stanford dataset, (ii) cross-resistance to the inhibitors within a drug class, (iii) if including context-dependent information (close positions within the sequence, as well as long-range/structural data) improves drug prediction, and (iv) assessing if the information given by these different kernels is redundant or complementary. For the rest of this work, HIV protease and PIs will be used as the case study.

## 2 Material and methods

### 2.1 Dataset and Preprocessing

All protease sequences were obtained from the PI dataset of the Genotype-Phenotype Stanford HIV Drug Resistance Database (version date: 2024-1-16). This dataset consists of 4504 isolates tested for their *in vitro* resistance to some (or all) of the following PIs: atazanavir (ATV), darunavir (DRV), fosamprenavir (FPV), indinavir (IDV), lopinavir (LPV), nelfinavir (NFV), saquinavir (SQV) and tipranavir (TPV). The resistance to each drug is quantified as the relative half maximal inhibitory concentration (*IC*50_*drug*_), defined as the ratio between the IC50 of the HIV isolate *in vitro* and the IC50 of the WT virus. Then, *IC*50_*drug*_ *>* 1 indicates that the isolate is more resistant to the drug than the WT virus. Following a common practice in the field, these ratios were log-transformed [Bonet, 2015]. Furthermore, resistance quantification in the Stanford Genotype-Phenotype dataset is done with several commercial and lab methods (usually Antivirogram or Phenosense). For the purpose of the present analysis, all entries using a method different to Phenosense were excluded.

Regarding the preprocessing of sequence data, the subtype B consensus protease was considered WT. All sequences longer or shorter than 99 amino acids, or with unknown residues (missing values or symbols like “X”, “B”, etc.) were removed. These events were uncommon: only 0.178% of all sequences had insertions, 0.755% truncations, and 15.6% missing or ambiguous values. Duplicated sequences coming from the same patient were removed to minimize bias. Also, there were 125 HIV samples of other subtypes than B (like HIV-1 C, G or U), which were reserved as an additional set. Thus, the final cleaned PI dataset consisted exclusively of HIV-1 B protease samples with missense mutations that came from a “Clinical” environment (as opposed to the “Laboratory” environment).

As complementary information, a crystal structure of WT protease was retrieved from the RCSB Protein Data Bank (PDB code: 3OXC). Euclidean distance (in Å) between all pairs of protein residues was computed from the (x,y,z) coordinates of their *α*-carbons.

### 2.2 Linkage disequilibrium

Linkage disequilibrium (LD) was computed for measuring the strength of non-random associations between residues in this dataset. Given two multi-allelic *loci A* and *B*, we can define the LD between any possible pair of alleles in these *loci* as:

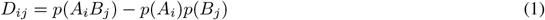

where *p*(*A*_*i*_) is the observed frequency of i-th allele at *locus A, p*(*B*_*j*_) is the observed frequency of j-th allele at *locus B*, and *p*(*A*_*i*_*B*_*j*_) is the observed frequency of *A*_*i*_*B*_*j*_. The correlation between *A*_*i*_ and *B*_*j*_ is then obtained as follows:

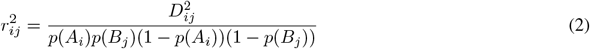

and then *r*^2^, the overall correlation between two *loci* (in our case: two protein positions), can be obtained following Zhao et al. [2007]:

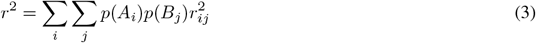

### 2.3 Kernel functions

Intuitively, a kernel can be understood as a two-place function that measures the similarity between two objects. In a slightly more formal definition, we could say that a given real symmetric two-place function is a kernel iff, for every finite set of objects, it generates a real-valued matrix that is squared, symmetric and positive semi-definite [Shawe-Taylor, 2004]. This kind of matrix is called “kernel matrix” and it stores the pairwise comparisons performed by the kernel function. Therefore, from then on, objects (in our case: the HIV sequences) are no longer represented explicitly. Different kernel functions have different criteria according to which they consider two objects more or less “similar”, but these criteria are not always obvious. To render them explicit, we should find the so-called “feature space” associated with the kernel. In fact, all kernel functions can be written as inner products in some feature space:

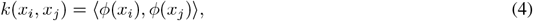

where *ϕ*(*x*) is the function that maps an object from the original (input) space to the feature space. Typically, the kernel function *k*(*x*_*i*_, *x*_*j*_) uses this mapping implicitly, ensuring that the process happens “under the hood”. This is known as “the kernel trick” and avoids the cost of explicitly mapping the dataset into feature space (which sometimes is *very* high dimensional) and then computing the inner product. Despite these advantages, in order to fully understand the criteria used by the kernels presented in this paper, we will make *ϕ*(*x*) explicit.

Before continuing, consider the following normalization:

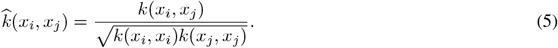

This is called the cosine normalization, which bounds the maximum possible value for a kernel to 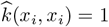. The feature space is normalized accordingly to have unit length:

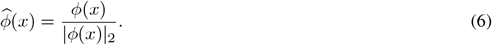

All kernels proposed in this paper are summarized at Table 1. Two kernel functions compare pairs of sequences with respect to their individual residues (similar to our approach in Ramon et al. [2019]), two work with *k*-mers and take into account close-range context, and the last two look for the interactions between residues, and thus can detect long-and close-range associations. Furthermore, three of these kernels take into account the amino acid mixtures, and their counterparts (the other three) do not.

**Table 1:** Kernel functions set-up.

|  | Without allelic mixtures | With allelic mixtures |
| --- | --- | --- |
| Amino acid substitution (Sequence) | Dirac kernel | Intersect kernel |
| Close-range context ($k$-mers) | Spectrum kernel | mixSpectrum kernel |
| Long and close-range context (pairwise interactions) | Interaction-Dirac kernel | Interaction-Intersect kernel |

#### 2.3.1 The Dirac kernel

The Overlap kernel is the simplest kernel function for categorical data. It assigns 1 if the two instances compared are equal and 0 otherwise:

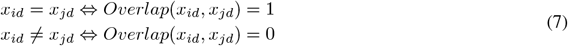

where *x*_*id*_, *x*_*jd*_ are the mutations or alleles in a given protein position (*d* = 1, 2,…, 99) in two protease isolates, *i* and *j. ϕ*_*Overlap*_(*x*) is the function that performs one-hot encoding. Therefore, in feature space, we find as many dummy variables as possible mutations in the *d*-th position. These dummy variables take a binary value (0 or 1) to indicate the absence or presence of a mutation. At this stage the mixtures are not taken into account: only one dummy variable per position can be 1.

For comparing full protease sequences, we can average the individual Overlap kernel’s evaluations along the *D=99* residues of the enzyme. This results in the Dirac kernel:

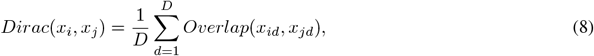

which is a valid kernel because it is a convex combination of kernels. The feature space for our Dirac kernel is obtained “concatenating” the feature spaces for all the 99 protease positions, and then scaling them by 1*/*99.

#### 2.3.2 The Intersect kernel

The Intersect kernel measures the similarity between two finite sets. It can be used to handle amino acid mixtures. For isolates *i* and *j*, let *X*_*i*_, *X*_*j*_ be the non-empty sets of alleles in the *d*-th position. Then:

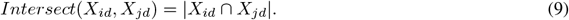

To remove the size effect and have a result bounded between 0 and 1 (like in the Overlap kernel), we can apply the cosine normalization:

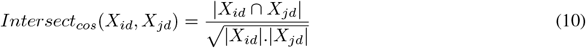

We can map our data onto feature space via multi-hot encoding. Thus, *ϕ*_*Intersect*_(*X*) recodes a set coming from a universe with *l* elements into *l* dummy variables (each one representing a given level) that can be 1, if a given element is present in the set *X*, or 0 if it is not. Then, it is trivial to see that, in absence of mixtures, this kernel is identical to the Overlap kernel.

As we did in Equation 8, we can average the individual Intersect kernel’s evaluations along the full protease sequence:

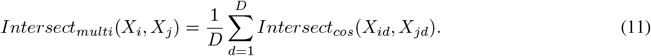

For simplicity, we will call this last kernel “the Intersect kernel” from now on. The feature space is also obtained as in the Dirac kernel.

#### 2.3.3 The Spectrum kernel

Let *A* be the 20 canonical amino acids, *x*_*i*_ be a protease sequence (a linear polymer of the amino acids defined in *A*), and *s* a *k*-mer of *x*_*i*_, *i*.*e*. a subsequence of length *k*. We can count the times (*t*_*s*_) that *s* occurs in *x*. If we do the same for all possible *k*-mers, each *t*_*s*_ becomes a feature in feature space, which has dimension |*A*|_*k*_.

The Spectrum kernel is the dot product of two strings mapped onto the feature space we have just defined:

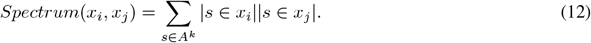

This kernel ranges from 0 to any positive number. Again, we can cosine-normalize the Spectrum kernel (Equation 5) so the result is bounded between 0 and 1.

In this work, the *k*-mers will have length *k* = 1, 2, 3. With *k* = 1, the Spectrum kernel compares sequences according to their amino acid compositions. With *k* = 2 and *k* = 3, the criterion is the frequency of each possible dimer and trimer, respectively. As these substrings consists of contiguous residues, this kernel looks for close-range context.

#### 2.3.4 The mixSpectrum kernel

This is a modification of the Spectrum kernel that can accept mixtures. Consider, for instance, a mixture in the *d*-th position of the *i*-th sample: *X*_*id*_. The number of amino acids in the mixture is |*X*_*id*_|. If *k* = 1, each one of the |*X*_*id*_| amino acids will contribute to its corresponding *t*_*s*_ with 1*/* |*X*_*id*_|. If *k* = 2 and two contiguous positions *X*_*i*(*d−*1)_, *X*_*id*_ contain mixtures, all possible dimers will be created: *i*.*e*. all possible combinations between both positions, so each one of them will count as *t*_*s*_ = 1*/*(|*X*_*i*(*d−*1)_||*X*_*id*_|). If *k* = 3, all combinations among the three positions will be performed, and so on. For instance, if *X*_*i*(*d−*1)_ = *A* and *X*_*id*_ = *LKR*, the *k*-mers spanning these two positions will be: {*AL*}, {*AK*}, {*AR*}, and they will be count as 1*/*3 in feature space. If the other position contains also a mixture, for instance *X*_*i*(*d−*1)_ = *AV*, this will result in: {*AL*}, {*AK*}, {*AR*}, {*V L*}, {*V K*}, {*V R*} with a weight of 1*/*(3 × 2). It is easy to see that, in the absence of mixtures, *t*_*s*_ = 1 and the kernel is equivalent to the Spectrum kernel.

In this work, the mixSpectrum kernel is cosine-normalized (see Equation 5), so the final result is bounded between 0 and 1.

#### 2.3.5 The Interaction kernel

Polato and Aiolli [2021] describe a kind of propositional kernel that counts the number of conjunctions formed using *d* binary variables in **x**_*i*_, **x**_*j*_ ∈ {0, 1 ^*P*^}. As our kernels for categorical variables implicitly map data into feature space using the one-hot encoding, we can relate them to the propositional kernels doing:

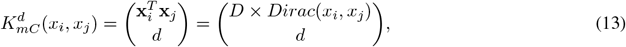

If the conjunctions we are interested in are pairwise interactions, then *d* = 2. The feature space consists of all possible interactions between two alleles at different positions. For instance, consider the feature *A*_*i*_ × *B*_*j*_, which denotes the conjunction between the *i*-th allele at position A and the *j*-th allele at position *B*. This feature will be 1 if a specific isolate has both *A*_*i*_ and *B*_*j*_, and 0 if it has only one or the other, or none.

For our HIV data, we are mostly interested in pairwise associations between positions that may be distant when taking into account the protease primary structure (sequence) but that are more or less “in the vicinity” when considering the folded enzyme. Thus, using the structural information of the 3OXC protease, only interactions between pairs ≤ 15 Å apart in the WT protease will be considered. Then, the Interaction kernel is computed following Equation 4. Finally, Equation 5 is applied so the resulting kernel is bounded between 0 and 1, just like the rest of kernels in this paper.

#### 2.3.6 The Intersect-Interaction kernel (Interaction kernel for mixtures)

Following the same reasoning than in Equation 13, we could expand the Interaction kernel to handle amino acid mixtures. However, such a kernel does not only take into account the pairwise interactions between sets (here: protease positions), but also the pairwise interactions between the elements inside a set (*i*.*e*. the different amino acids within a mixture). As here we are assuming strictly structural/functional associations, these “interactions within” are removed of the feature space, as well as those between residues that are farther than 15Å in the WT protease. The Interaction kernel for mixtures is then computed with Equation 4 and cosine-normalized with Equation 5.

### 2.4 Amino acid mixtures vs sampled sequences

Kernels that could handle mixtures and those that could not (see Table 1) were compared via three different metrics: Coinertia RV coefficient, Procrustes *r*, and Kendall’s *τ* coefficient. First, mixtures were sampled at random 30 times, each time keeping only one allele, and thus 30 “versions” of the PI dataset were presented to the Dirac, Spectral and Interaction kernels. The resulting 30 kernel matrices per kernel type were combined in a consensus matrix doing:

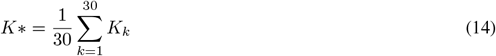

Then, the kernel Principal Components Analysis (kPCA) of this consensus matrix was computed and compared to the corresponding kPCA of the “kernel for mixtures” with the RV and Procrustes *r* coefficients. The approach was a bit different for computing the Kendall’s *τ* coefficient. In this case, to assess the dispersion of each one of the 30 “versions” of the dataset, 30 kPCAs were computed; then, the first Principal Component (PC1) of each one was contrasted to the PC1 obtained with their “kernel for mixtures” counterpart.

### 2.5 Training and test set-up

After pre-processing, the main dataset had 2087 HIV-1 subtype B isolates, while the additional dataset consisted of 125 samples of other subtypes. To assess the prediction models’ performance, the main dataset was split at random in training (80%) and test (20%) sets. To further ensure the independence of both sets, all samples coming from the same patient were always placed either in the training or in the test set. Only 243 patients in the dataset were in this situation, and each one of them contributed with 2 to 5 non-identical samples.

The three sets (training, test and additional test, amounting to 2212 sequences) were visualized together with kPCAs to see if they formed distinct clusters. To further assess if the additional test was significantly different from the main dataset, a kernel-based hypothesis testing for homogeneity was computed using the Intersect kernel. The test statistic was Maximum Mean Discrepancy (MMD).

After that, the training set was used to generate the prediction models, which were computed for each protease inhibitor separately. Thus, the training set size was different across models, as not all HIV isolates had been tested for all drugs. FPV training set had 1593 isolates, ATV 1228, IDV 1631, LPV 1481, NFV 1672, SQV 1639, TPV 990 and DRV 903.

A 5×2 nested cross validation set-up was used to do hyperparameter optimization (inner loop) coupled with the comparison of the different kernels used (outer loop). Hyperparameter optimization was done through grid search (candidate values: *C* = 0.1, 1, 10, 100, while *ϵ* = 0.1). Then, the final models for each drug and kernel were built using the whole training with the hyperparameters obtained with the 5×2 procedure. To assess the model performance in both the test set and the additional test, the NMSE (Normalized Mean Square Error) between actual (*y*) and predicted (*y’*) drug resistances was computed:

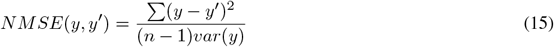

NMSE can be understood as the fraction of target variance not explained by the model, and it is related to the coefficient of determination *R*^2^:

### 2.6 Obtaining and comparing feature importance rankings

#### 2.6.1 kernel PCA

Principal Component Analysis (PCA) can be computed using Singular Value Decomposition (SVD). If **X** is a centered real-valued dataset (all columns have zero mean), **X** = **UΛV**^*T*^. Then, new variables (Principal Components or PCs) can be obtained in two ways:

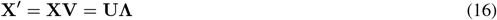

The former way projects the original dataset onto the **V**, the Principal Axes. The latter way is relevant to kPCA. The SVD of any kernel matrix is: **K** = **UΛ**^**2**^**U**, which (due to the kernel trick) allows to obtain the projection even when the original data is not real-valued. This by-passes **V**, as the Principal Axes now live in feature space. To recover this information (which allows us to compute the contribution of each original feature to the PCs), we need to revert to the first approach, which means computing the PCA in feature space.

#### 2.6.2 SVM

SVM is a supervised ML method that can be used for classification and regression. Consider for instance the following model:

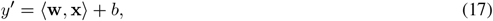

where **w** ∈ ℝ^*P*^ is a vector of weights, *b* the intercept, and *y’* is the prediction obtained from **x** ∈ ℝ^*P*^, a real-valued vector with *P* features. In our particular case, the *in vitro* resistance of a HIV isolate is *y* = log_10_*IC*50_*drug*_(*strain*), and **x** is the sequence information (point substitutions, *k*-mers, or interactions) in feature space. To obtain the importance of each feature we need **w**. Once we know the map *ϕ*_*k*_: *x* → **x** used implicitly by a particular kernel, we can map data into feature space and recover **w** in a rather straightforward manner:

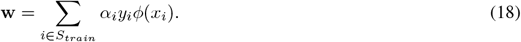

Here, *S*_*train*_ are the training samples and *α* the nonnegative Lagrange multipliers that weight each training sample. Contrarily to PCA, we do not need to map all data into feature space, only the Support Vectors. This is because Support Vectors are the only samples that contribute to the solution, as only they fulfill *α >* 0. In addition, we can observe that, due to the specific data and kernels used in this paper, all features in feature space are strictly nonnegative. Thus, the sign of **w** depends only of *y*, and we can deduce that a feature *p* contributes to drug resistance if **w**_*p*_ *>* 0 and, to the contrary, is related to drug susceptibility if **w**_*p*_ *<* 0. Finally, with the kernels we are using, features in feature space (dimension *P*) do not coincide with the original variables (dimension *D*). For instance, the Interaction kernel has as features the importances of pairwise interactions between all alleles at positions *A* and *B*. To obtain the overall importance of *locusA* × *locusB*, we can sum the (absolute) importances given to all those alleles.

In this work, the feature importances will be used to rank features from more to less relevant. Then, similarity between rankings will be measured with Kendall’s *τ* coefficient. Kendall’s *τ* coefficient, which by the way is a valid kernel, ranges between-1 and 1: 1 means that the two rankings are identical,-1 that one ranking is the reverse of the other, and 0 that the two rankings are completely independent.

### 2.7 Software

All the analyses were run in R. `kerntools` v1.2.0 was used to compute the kernel matrices and kPCAs, contribution of features to PCs, and SVM feature importances. The SVM models were trained with `kernlab` (v0.9-33), and the kernel-based hypothesis testing was computed with `rnaotai` (v0.2.5). The code for reproducing all experiments in this paper is available at https://github.corn/elies-rarnon/hivresistance/.

## 3 Results

Protease sequences in this dataset had high genetic diversity. The median number of mutations (with respect to the WT protease) was 9 (1st quartile: 5, 3rd quartile: 14). For the non-B subtypes, it was slightly higher: 10 (8.29, 11.23). Taking into account the three sets (training, test, and additional test), 58% of samples presented amino acid mixtures. The number of positions affected was 1 on average, but it was as high as 17 in some samples (see Figure 1 panel A). In total, 2168 sequences (out of 2212) were unique. All 99 protease residues, with the sole exception of P1 i W42, were polymorphic. Their range was 2-15 different alleles (amino acids) per position, with a median of 5. The most diverse positions were 63 (15 alleles), 12 and 37 (14 alleles), 34 and 72 (13 alleles), 19 (12 alleles), 10 and 67 (11), 20, 43, 48 and 89 (10). Excluding position 48, the remaining are considered minor or neutral mutations. Despite this large variability, the allelic distributions tended to be very skewed in favor of the major allele (Fig1 panel B), which was usually the WT (though not always). In general, the positions more prone to mixtures (10, 72, 37, 62, 71, 36, 63, 46, 13 and 77) had less skewed allelic distributions.

**Figure 1.**
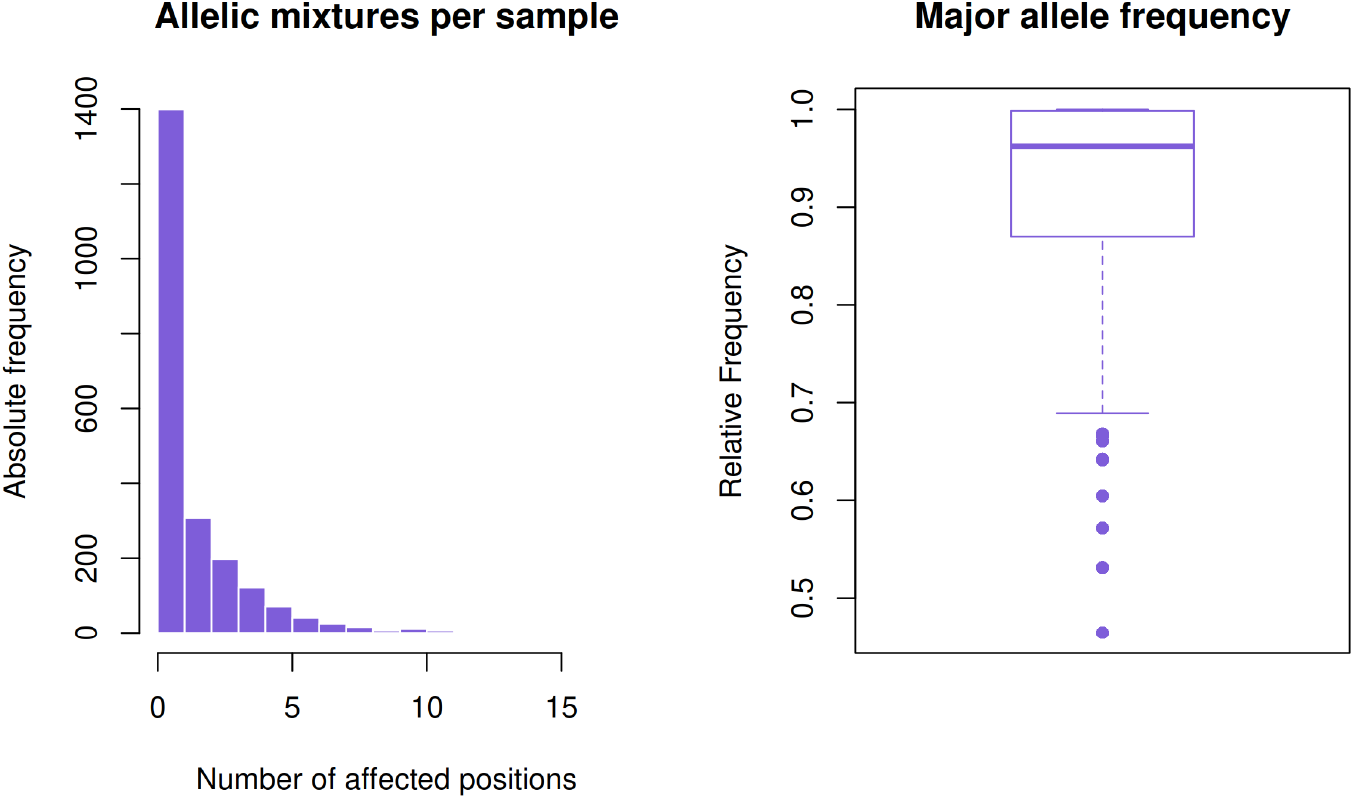
Protease diversity. Panel A: Number of allelic mixtures per sample. Panel B: Frequency of the major allele. The outliers, from bottom to top, correspond to the following protease residues: 10, 71, 36, 93, 46, 90, 54, 35, 82 and 62.

Protease LD is in Supplementary Figure 1. With some exceptions, pairwise correlation between positions is weak. Only 5 pairs had *r*^2^ ≥ 0.20: 30:88, 32:47, 20:36, 54:82 and 73:90. In this regard, Haq et al. [2009] also found 30:88, 32:47 and 54:82 to be the most correlated pairs, and observed that they involve spatially close residues. On average, the ten positions more strongly correlated to other positions were (from top to bottom) 54, 90, 71, 46, 33, 10, 32, 84, 20 and 73, which are also cited in previous literature, *e*.*g*. Wang and Lee [2007]. Three of them (10, 20 and 71) are minor positions, while the rest are major positions. This is consistent with the mechanisms followed by the protease to acquire resistance, where the conformational changes induced by major mutations are compensated by the minor mutations. Also, it agrees with previous literature that reports that, in treatment-experienced patients, LD in the protease/reverse transcriptase gene is mainly due to drug selection pressure.

Log-resistance distributions can be found at Supplementary Figure 2. The NFV, IDV and LPV distributions are bimodal in this dataset, with a relevant fraction of the isolates being resistant to these PIs. On the other end, TPV and DRV, the latest approved PIs [Wensing et al., 2010], have few resistant isolates. Both drugs impose a higher genetic barrier to resistance emergence than other PIs [Duffey et al., 2024, Ali et al., 2010]. This is the case indeed in this dataset; see Supplementary Figure 3 to see the logIC50 values as a function of the number of mutations for the eight PIs. Furthermore, they are both reported to inhibit protease dimerization, not only its enzymatic activity [Hayashi et al., 2014, Dakshinamoorthy et al., 2023].

### 3.1 Amino acid mixtures

All kernels that cannot handle mixtures in Table 1 have as a counterpart a kernel that can. To see to what extent they were divergent, the kPCAs computed from the kernels with mixtures were compared to those computed using the 30 resamples. To do so, three different measures were used (see Table 2). The first two (coinertia RV coefficient and Procrustes correlation) contrast PCA projections. According to these measures, the projections were almost identical (≥ 0.97). Next, the PCs were contrasted with Kendall’s *τ* coefficient. (The Interaction kernels were excluded because the high dimensionality of their feature space made very difficult to recover the PCs.) Here, to have an idea of the dispersion introduced by the sampling, the PC1 computed from the kernel for mixtures were compared to the 30 PC1s of the kernels over sampled data. Even then, the similarity was very high. For the case “Dirac vs Intersect” and “Spectrum vs mixSpectrum *k*=3”, Kendall’s *τ* matrix was visually displayed with a PCA (Supplementary Figure 4). The PC1 from the kernel with mixtures is placed on a central position, very close to the (0,0), while the 30 PC1s computed from the resamples without mixtures are scattered around it. In light of these results, only the “kernels for mixtures” were taken into account in subsequent experiments.

**Table 2:** Similarity between kernels computed from the dataset with mixtures and without mixtures (30 resamples).

| | Coinertia<br>RV coefficient | Procrustes<br>$r$ | PC1 correlation<br>median Kendall's $\tau$ (1st quartile, 3rd quartile) |
| --- | --- | --- | --- |
| Dirac vs Intersect kernels | 0.998 | 0.978 | 0.918 (0.915, 0.920) |
| Spectrum vs mixSpectrum kernels $k = 1$ | 0.998 | 0.970 | 0.979 (0.961, 0.979) |
| Spectrum vs mixSpectrum kernels $k = 2$ | 0.999 | 0.975 | 0.958 (0.956, 0.961) |
| Spectrum vs mixSpectrum kernels $k = 3$ | 0.999 | 0.980 | 0.896 (0.894, 0.898) |
| Interaction-Dirac vs Interaction-Intersect kernels | 0.998 | 0.988 | - |

### 3.2 Genotype vs phenotype ordination

kPCA was used to visually summarize the dataset. Figure 2 shows the kPCAs for the Intersect, Spectrum *k* = 2, Spectrum *k* = 3, and Interaction kernels. The corresponding Spectrum *k* = 1 plot is in Supplementary Figure 5 panel A. Dot color represents resistance to LPV; the plots of the remaining drugs are in Supplementary figures 6-12. In general, all kPCAs show a gradual but quite clear-cut separation between the resistant and susceptible isolates. This runs along the PC1 and, to a lesser extent, the PC2 (except for the Spectrum-3 kPCA). Only for DRV and TPV this spectrum-like separation seems less clear. It is also immediately visible that they are the PIs with less resistant samples (see Supplementary Figures 11 and 12). All in all, the kPCAs show that the main source of genetic variation in this data is related to resistance and also (because of the high concordance among the different protease inhibitors) cross-resistance. Both findings are consistent with data drawn from a clinical population.

**Figure 2.**
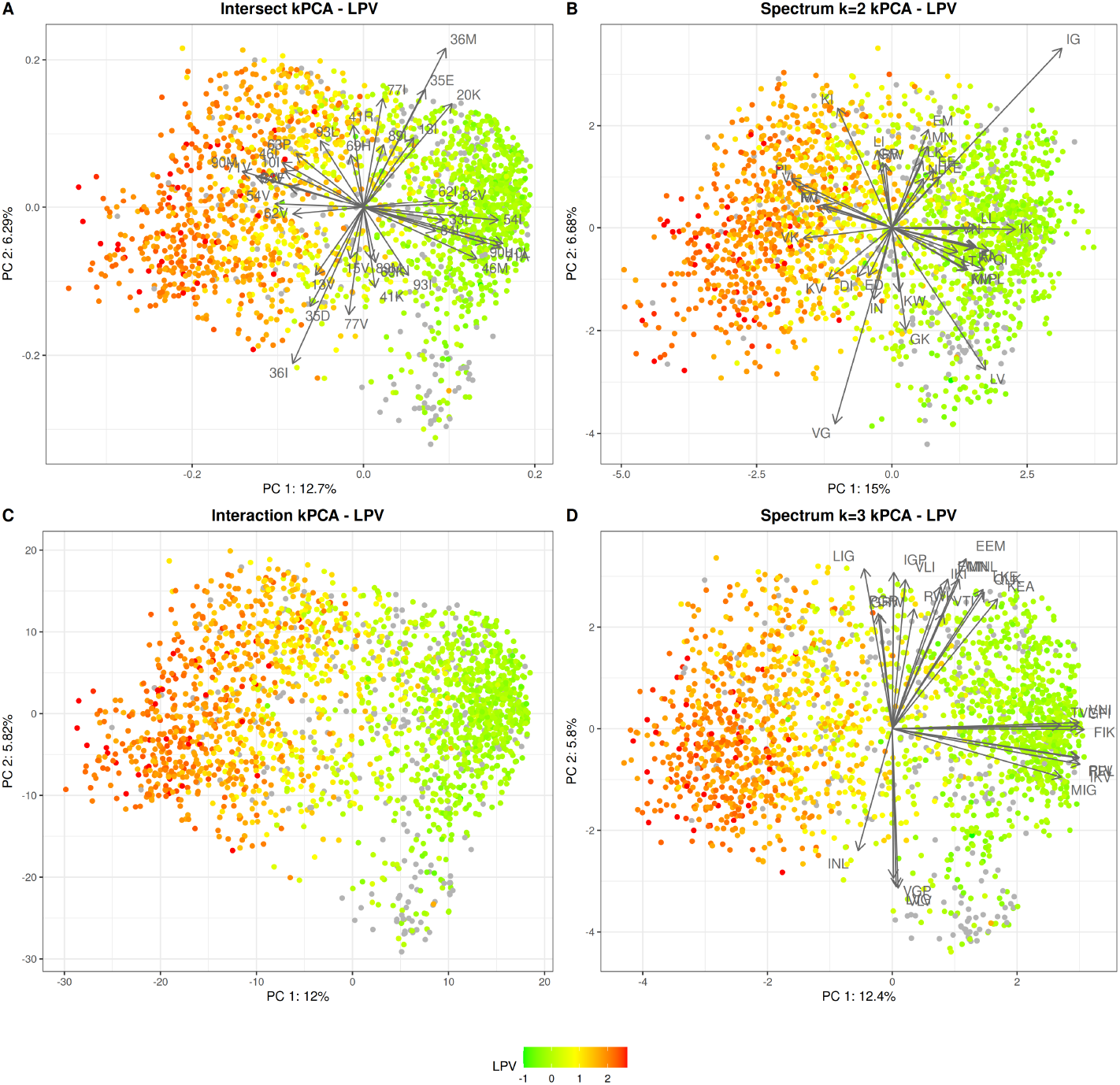
Genotype vs phenotype ordination. Panel A: Intersect kPCA, panel B: Spectrum-2 kPCA, panel C: Interaction kPCA, panel D: Spectrum-3 kPCA. Each dot is a protease isolate, colored according to its log-resistance value to LPV. Red means higher resistance values, while green means higher susceptibility. Sequences with missing IC50 data for the LPV drug are in gray.

Then, the kPCA colors were changed to highlight the training, test and additional test sets. As it can be seen in Supplementary Figure 13, there is a complete overlap between the training and test samples, which consist exclusively of subtype B samples; however, the additional test set samples (other subtypes) are clustered in the bottom right corner of the projection. That is, most variation captured by the PC2 is related to the genetic variation across HIV subtypes. To further verify that the additional test’s samples come from a different distribution, a kernel-based hypothesis test for homogeneity, based on the MDD statistic, was computed. The kernel used was the Intersect kernel. Differences between the test and additional test were statistically significant, as were differences between the training set and additional test: in both cases, p-value = 0.001. Instead, the training set vs test set scenario resulted in a p-value = 0.58.

In Figure 2, it can be observed that the kPCA projections obtained with the different kernels are pretty similar. The next step was to compare them with the Coinertia RV coefficient (Table 3). Intersect, Spectrum-3 and Interaction kPCAs were quite similar (≥ 0.90), while Spectrum-1’s was very different to the rest, followed by Spectrum-2’s. The Spectrum-1 kernel compares sequences according to their amino acid composition and seems to be the least informative of the kernels. It displayed a substantial overlap between resistant and susceptible isolates in Supplementary Figure 5 panel A and, also, it did not cluster apart the non-B samples (same figure, panel B).

**Table 3:**
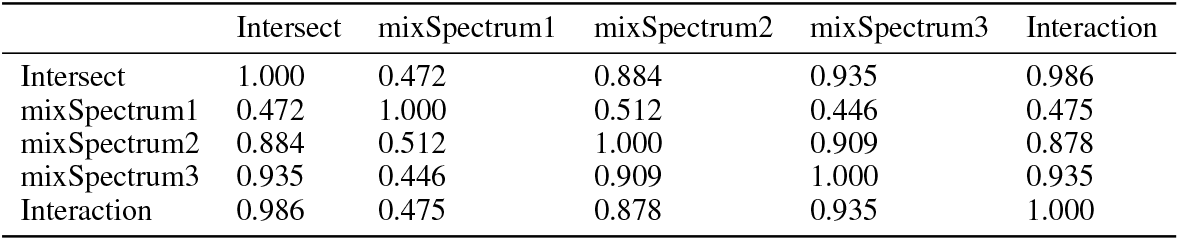
Pairwise similarity between sequence, *k*-mers and interaction kernels.

Next, the contribution of each variable (single residue allele/mutation, or *k*-mer) to the main PCs was studied. The most relevant are displayed as arrows in Figure 2 and in Supplementary Figure 14, which shows the top-20 variables with higher contribution. As expected, WT type alleles drive the projection to the right (more susceptibility), while mutated alleles drive it to the left (more resistance). Regarding the Intersect kPCA (panel A), it is immediately apparent that most WT type alleles associated with the low resistance area of the projection are paired with an alternative, mutated allele, driving samples to the high resistance area: A vs V at position 71, M vs I at 46, L vs I at 10, and so on. This is because, in feature space, dummy variables generated from one-hot (or even multi-hot) encoding have some degree of negative association. Furthermore, all mutations that are almost orthogonal to the resistance-susceptible boundary are previously described as resistant-related, either as major (L33F, M46I, I54V, V82A/F/T/S/L, I84V and L90M) or minor (L10I, I62V, L63P and A71V) [Wensing et al., 2022, Iyidogan and Anderson, 2014]. Not surprisingly, these positions also showed higher levels of LD in Supplementary Figure 1, and have less skewed allelic distributions compared to the average protease position (see Figure 1). Most of these residues have the WT allele as their 1st most frequent allele, a resistance-related alternative allele as the 2nd most frequent (thought in residue 63 the opposite is true), and then several other mutations that are much rarer.

Spectrum kPCA relevant *k*-mers (panels B-D) are more difficult to interpret. For ease of reference, results for Spectrum-3 kernel are summarized in Figure 3 panel A contrasted with those of the Intersect kernel. There, it can be observed that the trimers match around relevant single point amino acid mutations, especially at residues L10, M46, I54, A71, V82 and L90. Again, these positions are consistent with literature. VIG, the most relevant mutated motif, matches twice in the sequence (positions 71-73 and 84-86). Spectrum-2, which works with shorter *k*-mers, has more instances of multiple matches. That conveys some degree of ambiguity, especially because some positions are not described previously in literature. In any case, the majority of WT dimers in Supplementary Figure 14 panel B overlap with trimers in panel C (IK with FIK and IKV, PL and LV with PLV, KM and MI with KMI and MIG, KA and AI with KAI, LL, LT and TQ with LLT and LTQ). The IG motif is close to (or within) several important positions: 47-48, 50-51, 72-73, 85-86 and 93-94, but does not seem to play a great role in the context of the PC1. Regarding mutated dimers, PI seems related to RPI, VI to VIG, VK and KV to VKV and HKV, FV to GFV, MT to LMT and MTQ, and KI to PKI. Finally, IV as a mutated dimer matches positions 10-11 and 46-47 or 47-48.

**Figure 3.**
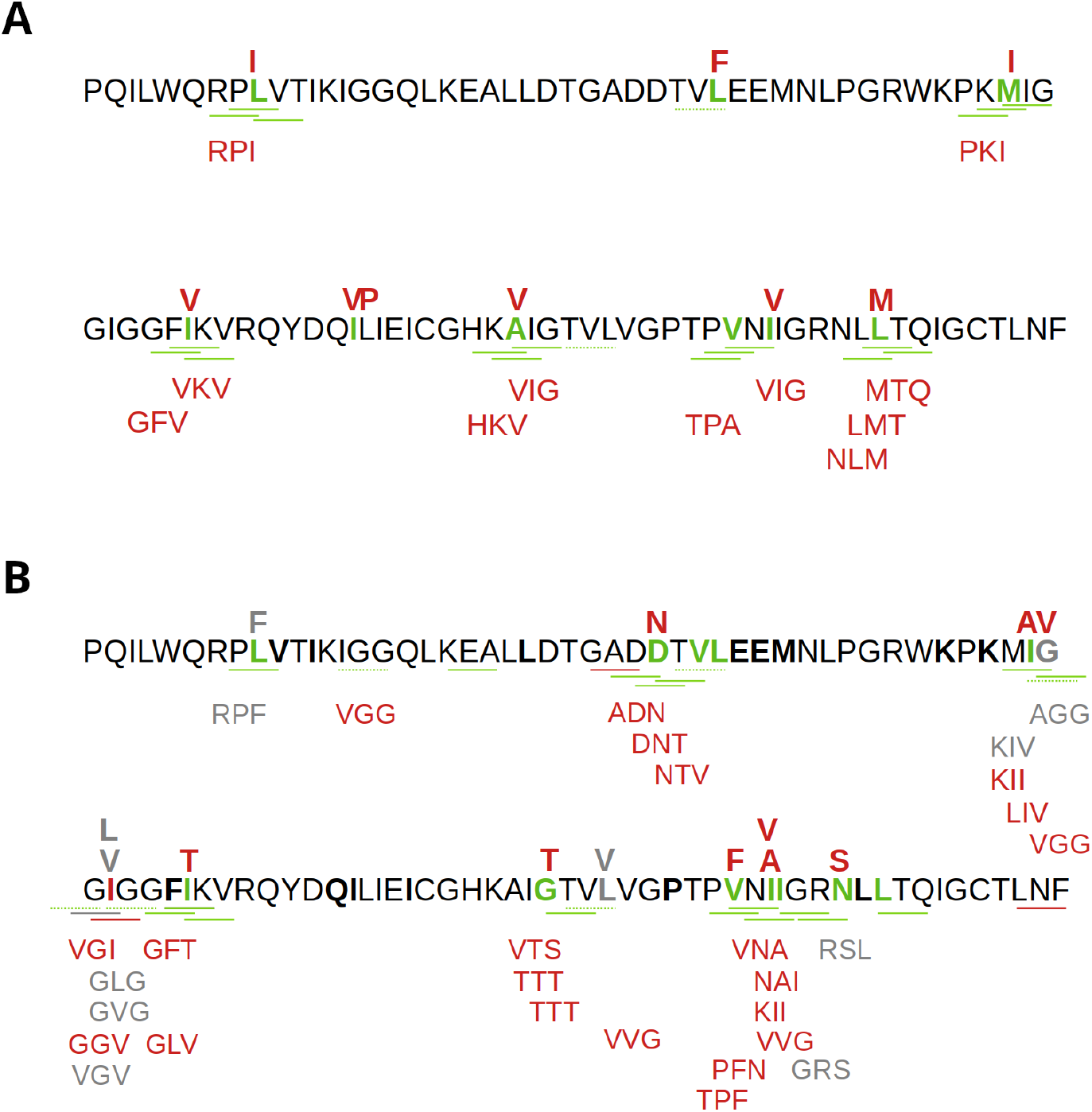
Consensus B protease sequence highlighting the most relevant alleles according to the Spectrum-3 and Intersect kernels kPCAs (panel A) and SVMs (panel B). Panel A: kPCA results. As Supplementary Figure 14 had only one mutated trimer, the present plot displays the following ten mutated trimers as well. Panel B: SVM results, as in Figure 5, but excluding: (i) non-WT trimers that increase susceptibility, (ii) non-WT that increase resistance to a single drug, except if related to mutation D30N. In both panels: WT key positions are highlighted in bold (for single amino acids detected with the Intersect kernel) or underlined (for trimers detected with the mixSpectrum k=3 kernel). Dotted lines underline trimers that match more than once. Single amino acid mutations are placed above the sequence, and mutated trimers are below. Red alleles are associated to higher resistance, green alleles to higher susceptibility, and grey alleles are associated to either one or the other depending on the drug.

As mentioned above, PC2 is related to subtype diversity (B vs non-B), and this is most clear in the case of the Spectrum-3 kPCA. The most contributing trimers to PC2 are in Supplementary Figure 14 panel D. Residues 75-79 (trimers VLV, LVG and VGP) seem more conserved in non-B subtypes. The Intersect kPCA points to a similar finding: in position 77, subtype B sequences are more prone to have the alternative allele I (instead of the WT allele V) than the rest of subtypes. Conversely, positions 34-38 (EEM, EMN and MNL) are more conserved in the B subtypes. In the Intersect kPCA, this seems to correspond to a lower frequency of E35D and M36I mutations, which is supported by [Nastri et al., 2023]. Both mutations may also play a minor role on resistance. Non-B subtypes also have higher frequency of the trimer GKW (residues 40-42), consistent with the R41K mutation in the intersect kPCA, which has been observed in a lot of non-B subtypes [Ali et al., 2010]. The same can be said of trimers spanning the positions 11-15, centered around the mutation I13V, or 18-22, as the residue 20 seems more prone to mutations in HIV-1 C and F than in B. Other relevant mutations detected by the intersect kPCAs are H69K (found also in HIV-1 C and F) and L89M (subtypes A, C and F) [Ali et al., 2010, Velazquez-Campoy et al., 2001, Sanches et al., 2007]). Lastly, some previous works have reported higher conservation of the WT alleles in residue 93 for subtype F, and 63 for C and F; see Nastri et al. [2023]. All in all, this is in line with previous knowledge that genetic variation among clades tends to be found outside the active site. Even so, these polymorphisms play a role in modulating enzymatic activity and increasing HIV replicative capacity [Ali et al., 2010].

### 3.3 SVM prediction models

#### 3.3.1 Performance overview

SVM point estimates of performance (validation, test and additional test) can be found at Supplementary Table 1. The final size of the test set varied among drugs, from 402 samples (NFV) to 215 (DRV). Figure 4 compares the bootstrapped NMSE in test (with error bars) and the 5×2 nested cross-validation NMSE. The Spectrum-1 SVM models performed significantly worse than the rest (NMSE between 0.37-0.70, depending on the drug). The other models are more even, although Spectrum-2 also had significantly worse performance in some protease inhibitors (for instance SQV). The general tendency was that the Intersect models performed slightly better than the *k*=2 and *k*=3 Spectrum models, while the Interaction models performed a bit better overall than the Intersect. Supplementary Figures 15 to 19 display the fitted vs actual log-resistance (in training) and predicted vs actual log-resistance (test) values. It can be observed that, in Spectrum-1 models, there was heteroscedasticity (more variance when predicting high resistance values) and a tendency to overfit. This reinforces that Spectrum-1 kernel, which compares the amino acid frequencies between isolates, is overly simplistic. Even then, the models generated with it had a certain degree of predictive power. There was also some amount of overfitting in the Interaction models and, to a smaller degree, in the Intersect models. In contrast, Spectrum-2 seemed more prone to underfitting. It should be remarked that the Interaction kernel generated feature spaces that were much higher dimensional than those of the rest of kernels (more than 13000 nonzero features vs less than 1000).

**Figure 4.**
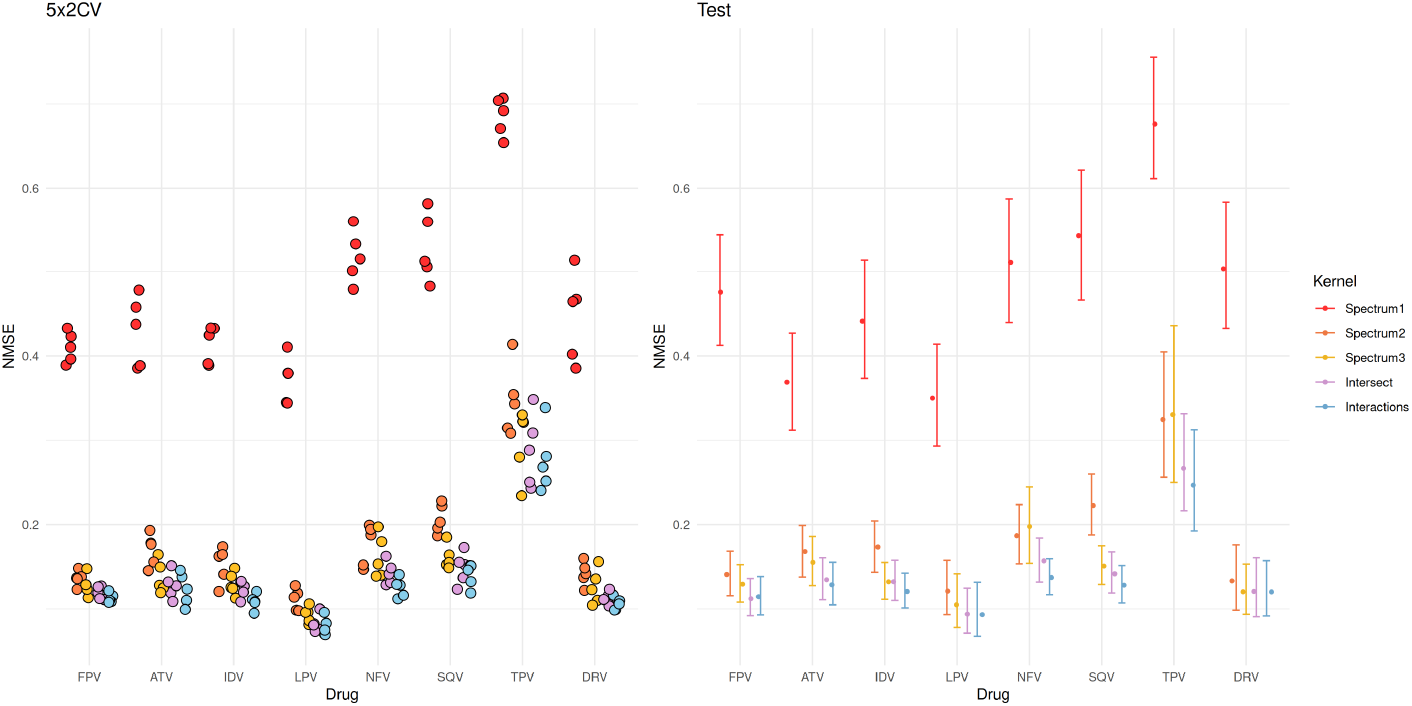
SVM performance. Panel A: 5×2 cross-validation error. Panel B: test error (estimates based on 3000 bootstrap samples) with a 95% CI.

If we focus on the best performing models, we can see (Figure 4) that the prediction error was low in all drugs (around 0.13), with the exception of TPV (NMSE ≈ 0.25). In this regard, other recent papers have also obtained significantly worse performances in TPV, *e*.*g*. Stolbova et al. [2024]. Intersect, Interaction and Spectrum-3 models had difficulties to correctly predict high resistance values of this PI (Supplementary Figures 15 to 19), though, as said previously, ther*e* are few isolates that are very resistant to TPV in this dataset. To a lesser extent, ATV had a similar problem regarding heteroscedasticity, despite having a lot more high-resistance examples for training.

In subsection 3.2, it has been shown that the test (subtype B samples) and the additional test (non-B samples) sets are significantly different. As a confirmation, performances of the SVM models (trained exclusively with B isolates) are much worse in the latter than in the former (Supplementary Table 1). Results should be interpreted with caution because the additional test set has a very low size and contains a mixture of non-B strains in different proportions; but this further suggests that models trained with subtype B protease sequences are not optimal for predicting non-B samples.

#### 3.3.2 Relevant alleles, mutations and interactions

For this section, only the feature importances of the best performing models (Spectrum-3, Intersect and Interaction SVMs) will be analyzed.

The 20 most important alleles and mutations according to each Intersect SVM model are in Table 4. Depending on the drug, these alleles explain between 17-21% of the total importance. As the training set size was different among the eight PIs, there may be some relevant but low-frequency alleles that are missing in some drugs. Amino acids associated to greater resistance as well as those associated to greater susceptibility are both compiled. The model balances both kinds of alleles, as the dummy variables in feature space are negatively correlated. For instance, some table entries only contain the WT allele (usually with a protective contribution). This means that mutations at this residue that increase resistance are placed lower in the ranking (*e*.*g*. L10, which has a plethora of resistance-associated mutations; see Wensing et al. [2022]). Oddly enough, there are few cases (like L76, I50, I54 in TPV and P79 in ATV) where the opposite is true: the WT allele seems to be detrimental. This indicates that some mutations make the virus more susceptible to a drug. For instance, it is well known that L76V increases susceptibility to ATV and SQV. Other mutations increase the resistance to several drugs but lead to a re-sensitization to others. Thus, I50L (specific to ATV) re-sensitizes to IDV, and N88S, a drug-resistance mutation against NFV, increases susceptibility towards FPV [Dakshinamoorthy et al., 2023, Bastys et al., 2020, Kozal, 2004]. In this vein, Schapiro et al. [2010] report that mutations L24I, I50L/V, I54L and L76V are related to an increased response to TPV, which agrees with the results presented here for these four protease positions.

**Table 4:** Top-20 alleles according to each SVM prediction model.

|  | FPV | ATV | IDV | LPV | NFV | SQV | TPV | DRV |
| --- | --- | --- | --- | --- | --- | --- | --- | --- |
| L10 |  |  |  | L |  |  | F |  |
| V11 |  |  |  |  |  |  |  | L |
| I13 | L |  |  |  |  |  |  |  |
| L24 |  | L | L | L |  | L | IM |  |
| D30 |  |  |  |  | DN |  |  |  |
| V32 |  | V |  |  |  |  |  | V |
| L33 | L |  |  |  |  |  |  | IL |
| E34 |  |  |  |  | A |  |  |  |
| E35 |  |  |  |  |  |  | G |  |
| M36 |  |  |  |  |  |  |  | T |
| K43 |  | I |  | Q |  | T | I |  |
| K45 |  |  | IK | R |  |  |  |  |
| I47 | AI |  | A | AI |  |  |  |  |
| G48 | ES | GV | EMV | EGV | V | GV | EG |  |
| I50 | LV | LV | IL | LV | IL | LV | I | LV |
| F53 |  |  |  |  |  | L |  |  |
| I54 | IM | IST | AILST | AIS | AST | IS | LT | IM |
| Q61* | G |  | D |  | G | G | D |  |
| I62 |  |  |  |  |  |  | M | M |
| I66* |  |  |  |  |  |  | V | V |
| G73 |  | GT |  |  |  | G |  | G |
| L76 |  | LV |  | V | L | L | L | LV |
| P79 |  | P |  |  |  | D |  |  |
| V82 | FT | S | FS | FLV | F | M | LT | FT |
| I84 | ACI | AI | AI | AI | ACIV | ACI | IV | IV |
| I85 | M |  |  |  |  |  | IL |  |
| N88 | DNS | NS | N |  | NS | DG |  |  |
| L89 |  |  |  |  | I |  |  | FL |
| L90 |  |  |  |  | L | L |  |  |
| Cum. imp.(%) | 17.7 | 19.1 | 19.7 | 19.1 | 20.9 | 21.3 | 16.7 | 17.9 |

Alleles/mutations highlighted in red contribute positively to drug resistance, while those in black contribute negatively. Positions that are usually considered neutral and/or absent of resistance catalogs are marked with an asterisk. Cum. imp. refers to the fraction of total importance explained by the top-20 mutations or alleles.

A major difference between the results presented in Table 4 and those of the Intersect kPCA (Figure 2) is that most mutations that SVM consider important have low frequencies. This allows to highlight some amino acid substitutions that eluded the Intersect kPCA but that are well-known in literature, especially D30N (NFV-exclusive), V32, I47A, G48V and I50L/V [Iyidogan and Anderson, 2014]. This aside, there is a good amount of concordance between both methods, particularly when they highlight positions 10 (LPV), 13 (FPV), 33 (FPV and DRV), 35 (TPV), 36 (DRV), 54, 82 and 84 (all PIs), and 90 (SQV and NFV), though SVM highlights rarer mutations. Also, at Table 4 there are some instances of positions rarely reported in literature (13, 35, 45, 79), or even considered neutral (61, 69). In return, it is striking the absence of the major position 46 (but not the adjacent residues) and the minor position 71. Both are downplayed by the SVM models and appear lower in the ranking: *e*.*g*. M46I is 36th for LPV, and the WT allele A71 is 24th for SQV). Nowadays, it is believed that M46I has a modest effect against PI binding, and that it plays a compensatory role instead [Bastys et al., 2020], as is the case of A71I, which has a stabilizing effect on the protease [Dakshinamoorthy et al., 2023].

The top-20 alleles for Spectrum-3 are in Figure 5 and (contrasted to those of Intersect kernel) in Figure 3 panel B. As in the kPCA analysis, relevant trimers tend to “swarm” around important single position mutations, namely positions 27-32 (pointing to D30N and, probably, mutations at residue 32), 46-56, 73-76 and 81-88, and to a lesser extent, 9-11 (L10), 71-76 (L76), and 90-92 (L90). In addition, some trimers include two or three amino acid substitutions; *e*.*g*. KIV, LIV, TTT, VTS (Figure 3 panel B). Mutation M46I is now acknowledged by several trimers, like KIV (which increases resistance to ATV and SQV) or MIG (more susceptibility to LPV). However, some amount of ambiguity remains, as there are trimers that match more than once (for instance, IGG has 3 matches, and LGG and TVL two each). In any case, structural elements around the active site are now perfectly recognizable: the protease flap region (residues 46–54), plus the two sides of the active pocket (residues 30-32 and 80-84) as well as nearby residues (85-90) some of which belong to the enzyme hydrophobic core. Similar hotspots have been observed by other authors, *e*.*g*. Stolbova et al. [2024], Su et al. [2016]). Stolbova et al. [2024] not only highlighted the subsequences 7–14, 31–35, 43–56, 67–74, 79–86, and 88–92, but also observed that they covered important resistance-associated mutations.

**Figure 5.**
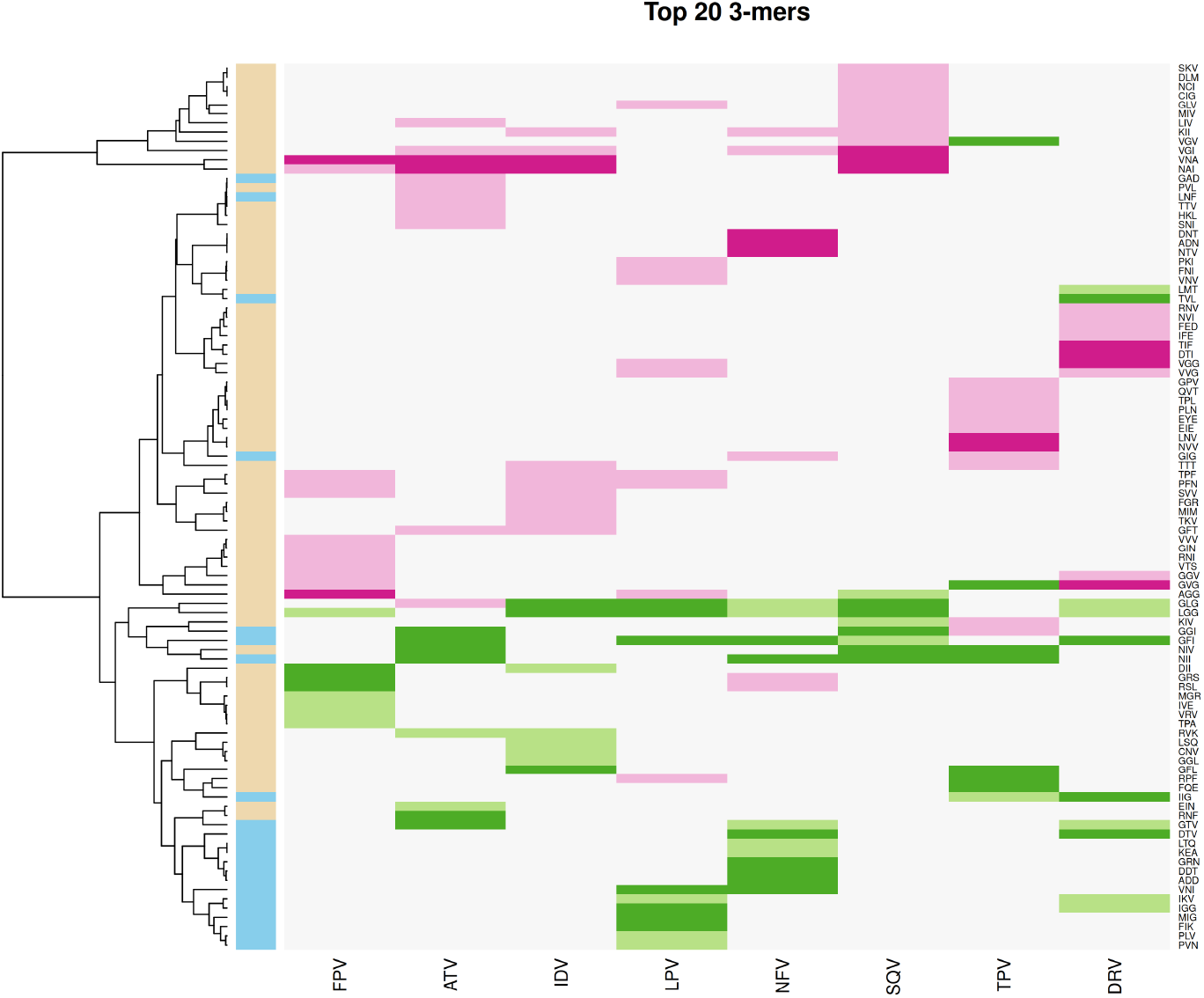
Spectrum *k* = 3 top 20 most important features (trimers). Stronger shades of pink denote a positive contribution to drug resistance, while green denotes a negative contribution. The left bar indicates if the trimer is found in the WT sequence (blue) or not (light brown).

As for the Interaction kernel, the top-20 interactions that affect drug resistance (according to SVM models) are at Table 5. Overall, ≃60% are interactions between resistance-associated positions (21 involving major×minor, 15 major× major, and 8 minor × minor), while the rest are mostly interactions between resistance-associated positions and neutral positions. There are also three pairs involving only neutral positions. The ten residues with more interactions on average are 54, 33, 35, 47, 48, 37, 10, 84, 82 and 83: 6 major, 3 minor and 1 neutral. Almost all important positions given by the Intersect kernels (Table 4) are present, with few exceptions, for example residues 30 or 90. On the other hand, very close-range interactions at the sequence level (35 × 37, 46 × 48, 47 × 48, 47 × 49, 71× 72, 82 ×83, 84 × 85 and 88 × 89) at Table 5 can be tied to some of the Spectrum-3 most important trimers discussed above. For instance, interactions I47× G48 and I47 × G49 are associated with LPV susceptibility, while I47A × G48 and I47A × G49 are related to LPV resistance; this corresponds to IGG vs AGG in Figure 5. Thus, interactions between I47 and the subsequent residues (*i*.*e*. the glycine-rich motive near the tip of the protease flap) seem relevant to resistance to this specific PI. EIN corresponds to E35 × N37. M46 × G48 is consistent with the protective trimer MIG (LPV), the double mutation M46L × G48V with LIV (SQV), and some other combinations like M46 × G48V and M46 × G48M with MIV and MIM. A71T × I72T is related to trimer TTT, and the double mutation V82F × N83N to PFN (more resistance to LPV and IDV). I84 × I85 increases susceptibility (trimer NII), in opposition to I84A × I85 (trimer NAI). Also, behavior of N88S × L89 (data not shown), related to FPV susceptibility and NFV resistance, is consistent with that of the trimer RSL. Despite this concordance, neither the Intersect nor the Spectrum-3 kernels highlighted so many residues that are usually considerated neutral. This phenomenon may arise due to overfitting (some interactions detected by the Interaction kernels may be not meaningful but caused by chance), although it should be noted that several works have reported covariation between neutral and resistance-associated mutations in HIV protease. Moreover, it has been shown in previous literature that some associations that occur in therapy-naive isolates are lost or reversed under the positive selection imposed by PIs, but not all [Hoffman et al., 2003, Rhee et al., 2007, Wang and Lee, 2007, Haq et al., 2009]. These lingering covariation patterns suggest that certain residues, despite not being directly associated to drug resistance, play a role in general compensatory mechanisms [Hoffman et al., 2003]. Some of these positions are highly polymorphic with or without drug treatment and/or have unusually balanced allelic distributions (some examples are residues 12, 19, 35 or 37, see again Hoffman et al. [2003]). In the Interaction kernel models, the general trend is that WT × WT major× major or major × minor interactions in Table 5 lead to a drop in resistance, with very few exceptions. However, when the pair involves neutral positions, almost half of WT×WT interactions seem to increase resistance, while certain combinations of polymorphic mutations make HIV more susceptible. This seems to suggest that the former have a direct impact on drug resistance, while the latter may be more unspecific: for instance, playing a role on catalytic efficiency or viral fitness.

**Table 5:**
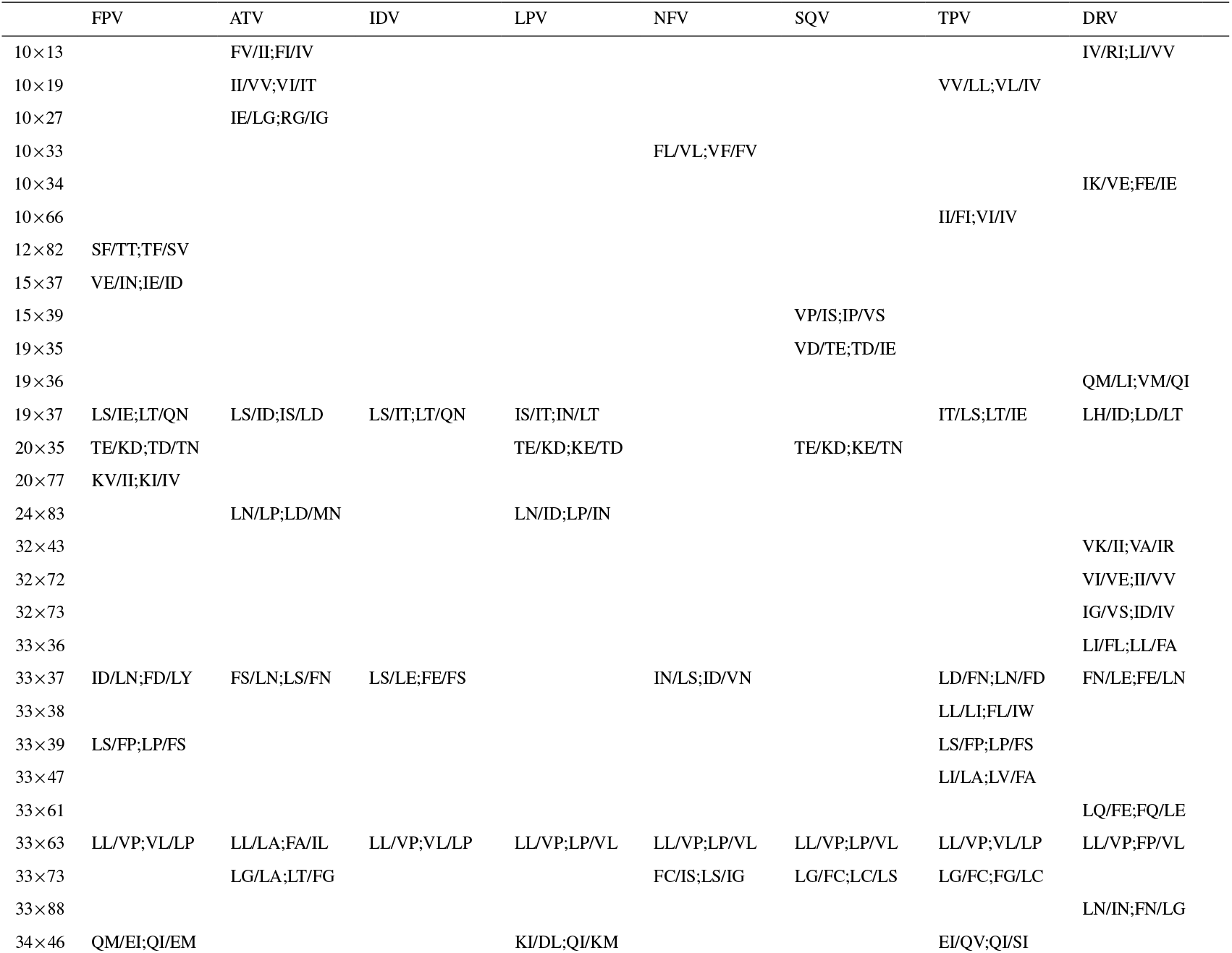
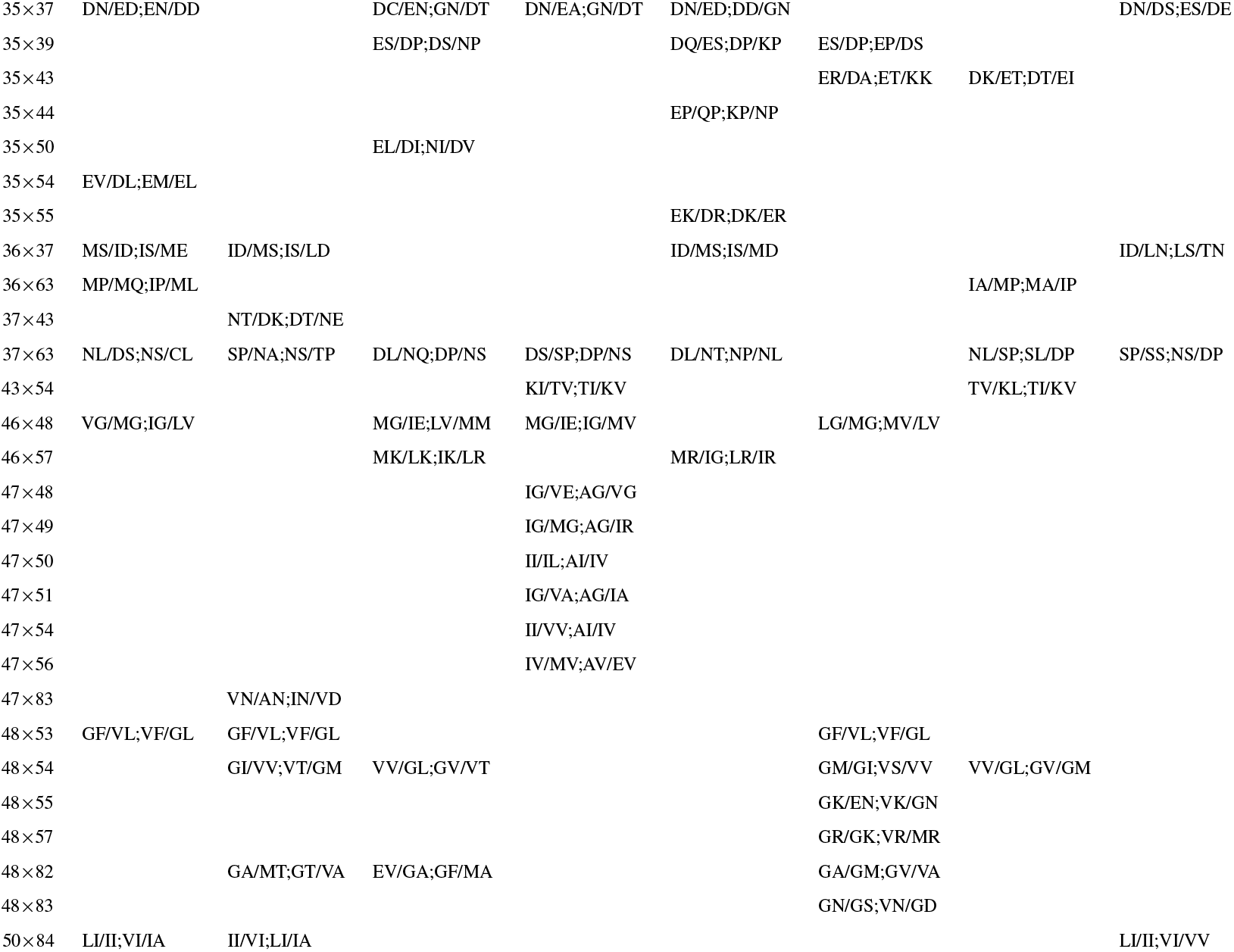
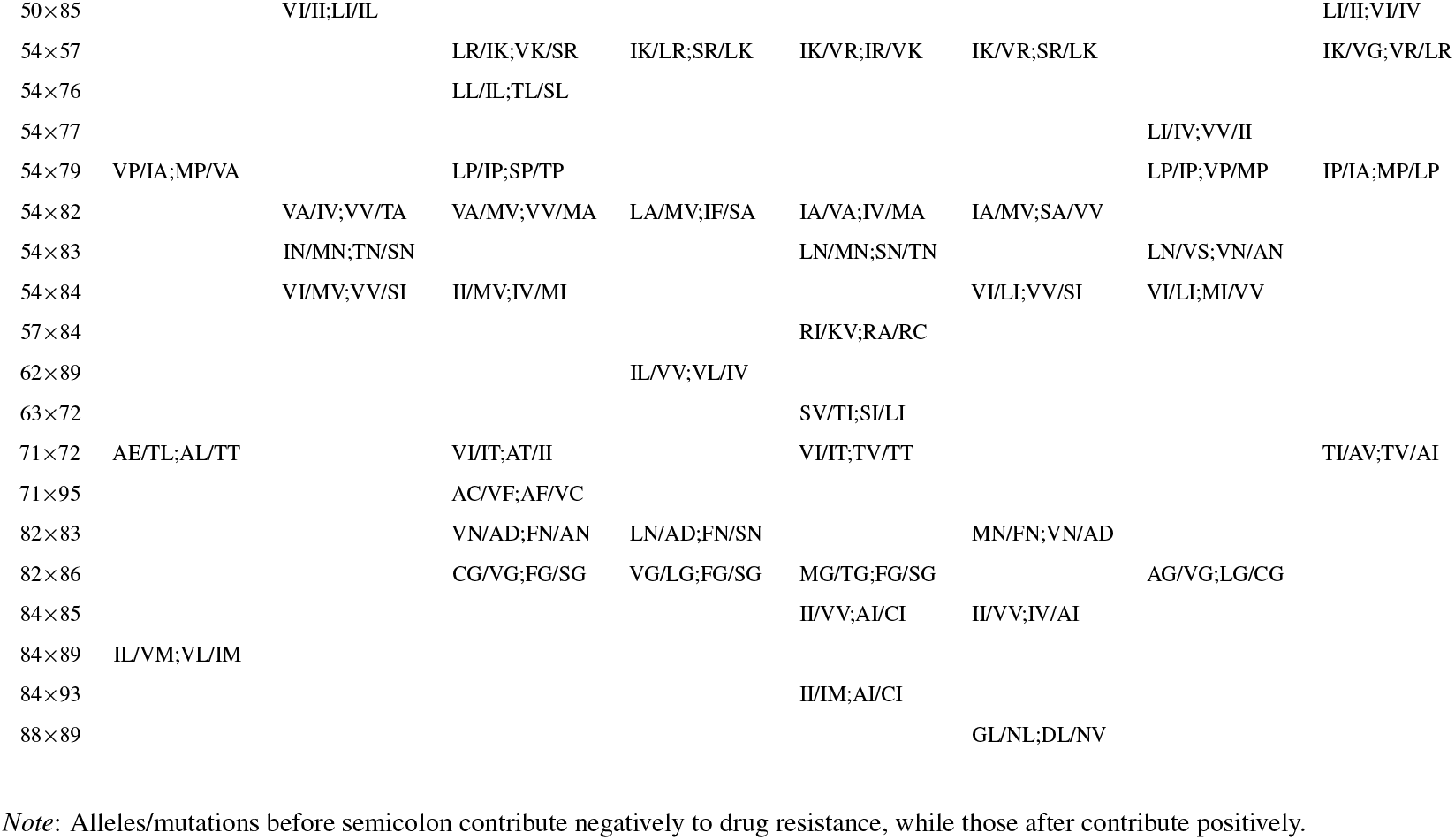
Top-20 interactions according to each SVM prediction model.

In this regard, it is intriguing the quantity of interactions found within the flap hinge loop. Segment 33-37 of the hinge domain is far away of the active site, but it is nonetheless highly polymorphic (in B as well as non-B subtypes), with residues having relatively common alternative alleles. In Table 5, positions 33, 35, 36 and 37 and 63 have relevant interactions with each other. 33 × 63 involves two of the three main sites of protease autoproteolysis. It has been proposed that mutations like L63P and L33F may stabilize the protease and/or protect it against autocatalysis [Yin et al., 2006]. According to the Interaction SVMs, WT × WT and L33V × L63P are (re)sensitize towards several drugs, as opposed to FP/FA/IA/IP. On the other hand, E35 regulates the opening and closing of the protease flaps [Tabler et al., 2024]; mutation E35D induces the curling of flaps, which may have an impact on inhibitor binding [Dakshinamoorthy et al., 2023]. Furthermore, E35D × N37D is reported by Hoffman et al. [2003] and Rhee et al. [2007], who note that, because the former mutation usually precedes the latter, it is likely that N37D is stabilized by E35D. Tabler et al. [2024] found that some combination of mutations at 35-37 (*i*.*e*. E35D × M36I × N37S) promoted premature protease activation, which inhibited particle production and increased the risk of producing defective viria, and Huang et al. [2011], that double mutants at residues 33 and 37 changed protease cleavage preferences, but single mutants did not.

Returning to the major residues, 48, 54 and 82 have relevant interactions with each other in Table 5 that, moreover, have been studied in previous literature. According to Hoffman et al. [2003] and Rhee et al. [2007], the three of them not only covariate, but they have a hierarchy. 82 tend to be mutated first, usually followed by substitutions at 54, while the mutations at residue 48 are likely to appear later. Unfortunately, this temporal component cannot be assessed with the Interaction SVMs. Besides that, mutation G48V, which is typical of SQV, causes steric clashes with the binding pocket residues and the PI, leading the flap to slide off and exposing the hydrophobic F53 to the solvent. This interaction is also in Table 5.

Lastly, as mentioned before, the pair with highest LD was 30:88, which are structurally very close. Unlike 54:82 (which we have just reviewed), 30:88 is absent of Table 5, even if D30N is a relevant mutation against NFV, causing a loss of catalytic efficiency that tends to be “compensated” by N88D [Haq et al., 2009, Hoffman et al., 2003]. In addition, N88S also interferes with NFV but via a different mechanism: changing the orientation of D30 [Dakshinamoorthy et al., 2023]. Altought this pair is not at Table 5, this is confirmed by the Interaction SVM results. Key interactions between these two positions are among the 1% percentile rank for NFV feature importances (data not shown). The double mutant D30N×N88D provided the highest increase on resistance to NFV, followed by D30×N88S. On the other hand, D30N×N88 increases most the susceptibility, followed by the WT×WT D30×N88.

#### 3.3.3 Cross-resistance

As seen in Table 4, the protease residues more related to cross-resistance were 50, 54, 82, 84 (all PIs), 48 (all PIs except DRV) and 88 (all excepting LPV, DRV and TPV). Putting aside the latter, all these positions are around the active site and the flap region. Mutations at 82 and 84 may have direct van der Waals contacts with PIs, and they are also related to expanded active site cavity and changes in flap position; this makes easier for the protease to release the PI [Dakshinamoorthy et al., 2023]. I50 is a key site within the flap and plays a crucial role in protease dimerization, as well as stabilizing the protease’s open conformation. I50V is attributed to the curling of flaps. Finally, 88 is distal to the active site but affect PI binding via its interaction with other positions, like 30 [Bastys et al., 2020]. It should be remarked that positions 54, 82 and 84 were already highlighted by the Intersect kPCA. They present more balanced allelic distributions, while 50, 84 and 48 are very skewed in favor of the WT allele. Mutations I84A/C/V (8 drugs), V82F/S/T (7 drugs), I50L/V, I54A/S/T and G48V (5 drugs each) are the most widespread according to Table 4. In general this is consistent with previous reports [Wensing et al., 2022, Iyidogan and Anderson, 2014]. Moreover, overall results at the same table further reinforce that TPV resistance profile is a bit different than those of other PIs. This has also been observed in previous works. For instance, Dakshinamoorthy et al. [2023] comment that TPV is the only nonpeptidomimetic PI, that it inhibits viruses resistant to most other PIs and that it binds directly to the I50 in the flap domain. Schapiro et al. [2010] say that mutations L24I, I50L/V, I54L and L76V, selected by other PIs, predicted increased response to TPV, which is consistent with Table 4, Lv et al. [2015] remark that TPV has a nonconventional HIV-1 protease resistance profile, and Boucher et al. [2018] that it retains activity against many LPV-and DRV-resistant viruses, having a role in salvage therapy. Even so, TPV is affected by some mutations at the cross-resistance residues I54, V82 and I84 (see also Wensing et al. [2010]).

The cross-resistant trimers were, with fours drugs each, VGI (positions 48-50), VNA (82-84), and NAI (83-85), and with three drugs each, KII (45-47 and 83-85), TPF (80-82) and PFN (81-83) (Figure 5). Instead, the protective trimers against resistance were LGG (15-17, 50-52) in six drugs, GLG (49-51) in five (and, as expected, it was resistance-related in ATV), GFI (52-54) in five, and NII and NIV (83-85) in four and three drugs each, respectively. Leaving out position 88, this follows closely the cross-resistance pattern explained above for the Intersect kernel.

Regarding the most widespread interactions (Table 5), 33 × 63 was relevant in all drug models, followed by other pairs at the hinge region like 37 × 63 (7 drugs), 33 × 37 (6 drugs) or 35 × 37 (5 drugs). This further supports that part of the perceived resistance modelled by the SVMs may be caused via an indirect or inespecific mechanism (*e*.*g*. affecting the viral fitness). Interactions involving major positions were also prominent, for instance 54 × 82 (5 drugs), 48 × 54, 46 ×48 and 54 × 84 (4 drugs each). The relevance given by the Interaction kernel to pairs that include cross-resistance residues like 54, 82 or 84 is consistent with the empirical finding that multi-drug mutants (which accumulate several major mutations stabilized with minor mutations) present expanded active-site cavities. As stated before, this weakens the binding of the PI to the protease, which can much more easily release it and continue to function in the interest of the virus [Dakshinamoorthy et al., 2023].

Finally, Kendall’s *τ* coefficient was used to compare the full feature importance rankings for the eight PIs. Supplementary Figure 20 confirms that TPV has the most different resistance profile, with DRV and ATV/SQV at the other two extremes of the spectrum. The singular traits of TPV may explain why the prediction models performed significantly worse that in the rest of drugs. FPV is at the same side than DRV (interestingly, FPV is the prodrug of amprenavir, which is very similar to DRV at the molecular level Lv et al. [2015]), while IDV, NFV and SQV are closer to ATV. Supplementary Figure 20 is similar to Su et al. [2016] Figure 1, who also commented that NFV, SQV, IDV (and LPV) are closely related, and so the likelihood that a protease mutation causes cross-resistance within this group is greater than for the other PIs Su et al. [2016].

## 4 Discussion

HIV sequence datasets are unusual when compared with datasets from DNA-based organisms. HIV protease has very few conserved residues, a wide range of alternative alleles (many of them in very low frequencies), and allelic mixtures caused by the quasispecies phenomenon. Also, most protease isolates in the Stanford Genotype-Phenotype Dataset are unique. This extremely high diversity makes HIV data difficult to tackle, posing several challenges to both classic rule-based studies and ML modeling. Preprocessing strategies like rare variant filtering, or sampling mixtures to keep only one allele, can mitigate the problem but downplay the emergence of novel mutations, which is an important mechanism for HIV to evade the antiretroviral therapy [Ramon et al., 2019]. Even then, ML algorithms have achieved great results in predicting HIV drug resistance. In particular, SVM is an effective and quite standard method for this particular problem [Stolbova et al., 2024, Dakshinamoorthy et al., 2023], that has been used for years (see Beerenwinkel et al. [2003]). However, some of the benefits it entails as a kernel method are not often capitalized, *i*.*e*. kernels’ ability to adapt to specific data and, in some cases, minimizing preprocessing. Therefore, kernels different to the “default” (the linear or the RBF kernels) continue to be underused.

The present paper proposes simple kernel functions that can effortlessly handle data with so much intrinsic variability: the Intersect, Spectrum, and Interaction kernels, all of them implemented in the R package `kerntools`. Unlike the Jaccard kernel proposed in Ramon et al. [2019], they are linear. Even if direct comparisons are not straightforward (because the Jaccard kernel was trained in an older version of the database, which included the non-B sequences), the kernels presented here reach similar performances in protease data. Moreover, it has been shown throughout this work that these linear kernels (despite being “nonconventional”) are transparent with regard to the criteria they use to grasp data (subsection 2.3). They also generate SVM models (and kPCA projections) that are explainable and interpretable, which is increasingly important in the medical field. In subsection 2.6, it is shown how the features they use and the weights they assign to them can be recovered from feature space. In contrast, obtaining (and interpreting) the feature importances of nonlinear kernels is possible but far more difficult (see for instance Cotter et al. [2011]). Studying the feature importances has contributed to: (i) compare SVM and kPCA to each other, (ii) contrast the feature importances with previously reported mutations and mutational patterns, and (iii) draw comparisons among the different kernels. In this respect, the main findings of this paper are:

i. Different kernel methods are complementary. In Ramon et al. [2019], keeping kPCA results side-to-side with those of SVM enhanced our understanding of SVM behavior, because they gave us an idea on how different kernels “saw” data. In the present work, there is indeed some amount of overlap between the kPCA and SVM results. However, kPCA stands on its own and provides an unique view of data (subsection 3.2). We can say that kPCA is primarily concerned with the main trends on genetic diversity that structure the dataset, while the goal for SVM is predicting a given target (here: drug resistance) as well as possible. By studying the first two PCs, it can be seen that kPCA spotlights *loci* (*e*.*g*. residues or *k*-mers) that have balanced (in the context of the HIV protease) allelic distributions and with some degree of LD among them. Not surprisingly, most of the leading alleles according to kPCA are known to cause cross-resistance. Secondarily, it also highlighted genetic differences between B and non-B HIV subtypes. With a dataset consisting of drug-naive HIV isolates, there is no doubt that the kPCA would paint a different picture; instead, when dealing with treatment-experienced patients, kPCA summarizes the most widespread alleles related to drug resistance in clinical populations. In contrast, SVM was able to focus on much lower frequency mutations. This attention to detail is essential because of the continuous emergence of novel polymorphisms in HIV, and the key role that rare mutations can play in resistance.
ii. The SVM models and kPCA are sound and consistent with previous literature. This has been assessed analyzing their top most important features and finding that, for the most part, they were already compiled in mutation catalogs like Wensing et al. [2022], Iyidogan and Anderson [2014]. Having interpretable algorithms that agree with previous knowledge is central to make ML models more trustworthy. Furthermore, it shows that kernel functions capture true biological meaning. This may push forward further studies about the role of specific/rare mutations or patterns on the enzyme conformation and function.
iii. Six different kernels that look upon data using different criteria have been studied (see Table 1). In a first step, this paper compared the impact of preprocessing the mixtures (keeping a single allele at random) onto the kernels. As multiple HIV variants coexist within a host as quasispecies, filtering or sampling the amino acid mixtures can distort the original data, especially if it is done a single time. Instead, the present work shows that sampling a dataset multiple times (*e*.*g*. 30 times) under the discrete uniform distribution and then averaging the kernel matrices approximates the behavior of the “kernels for mixtures” (Table 2). However, the “kernels for mixtures” are preferable because: (i) they are faster to compute (do not need additional pre-processing), (ii) require less memory space, and (iii) give a more accurate representation of the original dataset. In subsequent experiments, three kernels with different scope (point mutations, close-range context, and interaction between residues) were compared. The results showed that, in general, the three approaches present a high amount of redundancy when applied to the Stanford protease data (Table 3 and Figure 4). Among them, maybe the Spectrum kernels for *k*-mers were the least useful, especially when setting a very low *k*. In this regard, shorter *k*-mers have higher probability to match multiple times against a sequence. This obscures the interpretation and (worse still) can undermine the model if some matches are related to higher resistance and others to low resistance or neutral residues. The Spectrum kernel with *k* = 1 resulted in a very simplistic representation, and with *k* = 2 it generated models with a tendency to underfit data. *k* = 3 gave sounder models and some interesting insights into some drug-related mutational patterns, but had similar or slightly worse performance than both the Intersect and Interaction kernels. It is likely that the Spectrum kernel is better suited for problems involving sequence alignment, or longer sequences with a strong genetic linkage, so the close context holds a greater importance. The Intersect kernel is the most straightforward of the three, and it generated models that were very easy to understand, as they could be directly contrasted to previously published mutation catalogs. In comparison, the Interaction kernels suffered from greater complexity (even taking into account only pairwise interactions), with only a modest improving of performance.

The large number of different mutations in protease increases dramatically the potential interactions, while at the same time makes each specific combination very rare. The training set size is one order of magnitude lower than the feature space dimension, and there are signs of overfitting, which may cause these models to give weight to some interactions that are in fact caused by chance. Although additive effects may be have far greater weight that interactions, it is possible that in the future, with larger datasets, the Interaction kernel performance is improved. That said, it is remarkable the great similarity of the Intersect and the Interaction kernels matrices. A possible hypothesis is that this is a consequence of the entrenchment phenomenon. Protease residues co-evolve to escape the drug selective pressure, so in spite of the strikingly high mutation and recombination rates of HIV, the number of viable HIV variants is relatively small compared to all the potential variants [Wensing et al., 2010]. Isolates that carry only one or two major drug-resistance mutations are difficult to observe, because of the functional and structural constraints of the HIV protease, which is an enzyme under strong purifying selection Zhang et al. [2020]. Instead, they become entrenched because of the collective effort of the entire sequence (including mutations at neutral residues) to restore viral fitness. In other words, the Stanford HIV isolates used to train the Intersect and the Interaction kernels does not reflect all the (theoretical) protease mutational landscape; only a subset that, in addition, is already under strong epistatic interactions.

## 5 Conclusions

HIV has an extremely high genetic variability that hampers the study of resistance-associated mutations and patterns on a case-by-case basis, while also poses challenges to ML predictive models. In that context, kernel methods like SVM and kPCA are simple, reliable and transparent approaches for assessing drug resistance in HIV. This combination of good performance and interpretability is key in the current medical field. In particular, SVM reaches a very good prediction performance, and kPCA can be used to grasp the main genetic diversity patterns in data and see if they are related to resistance. Moreover, these methods can look at data through different prisms by changing the kernel function. In this work, three different approaches to analyze sequences have been used: single residue mutations, *k*-mers (close-range information in a sequence), and pairwise interactions (long and close-range information). It has been shown that, even when working with those “nonconventional” kernels, the models they generate can remain completely open and interpretable. The results presented here verify that these models are sound according to previous literature. Moreover, a thorough comparison among the three kernels has been done. In protease data, the close-range approach is maybe the least useful, and the most simple approach (point mutations) remains competitive.

## Declarations

### Availability of data and materials

The raw datasets are from the Genotype-Phenotype Stanford HIV Drug Resistance Database repository (https://hivdb.stanford.edu/pages/genopheno.dataset.htrnl). Crystal structure of WT protease can be found at: https://www.rcsb.org/structure/3oxc. Code used in this manuscript is available at https://github.corn/elies-rarnon/hivresistance/.

### Ethics approval and consent to participate

Not applicable.

### Consent for publication

Not applicable.

