## Supplementary Material for "Unraveling HIV protease drug resistance and genetic diversity with kernel methods"

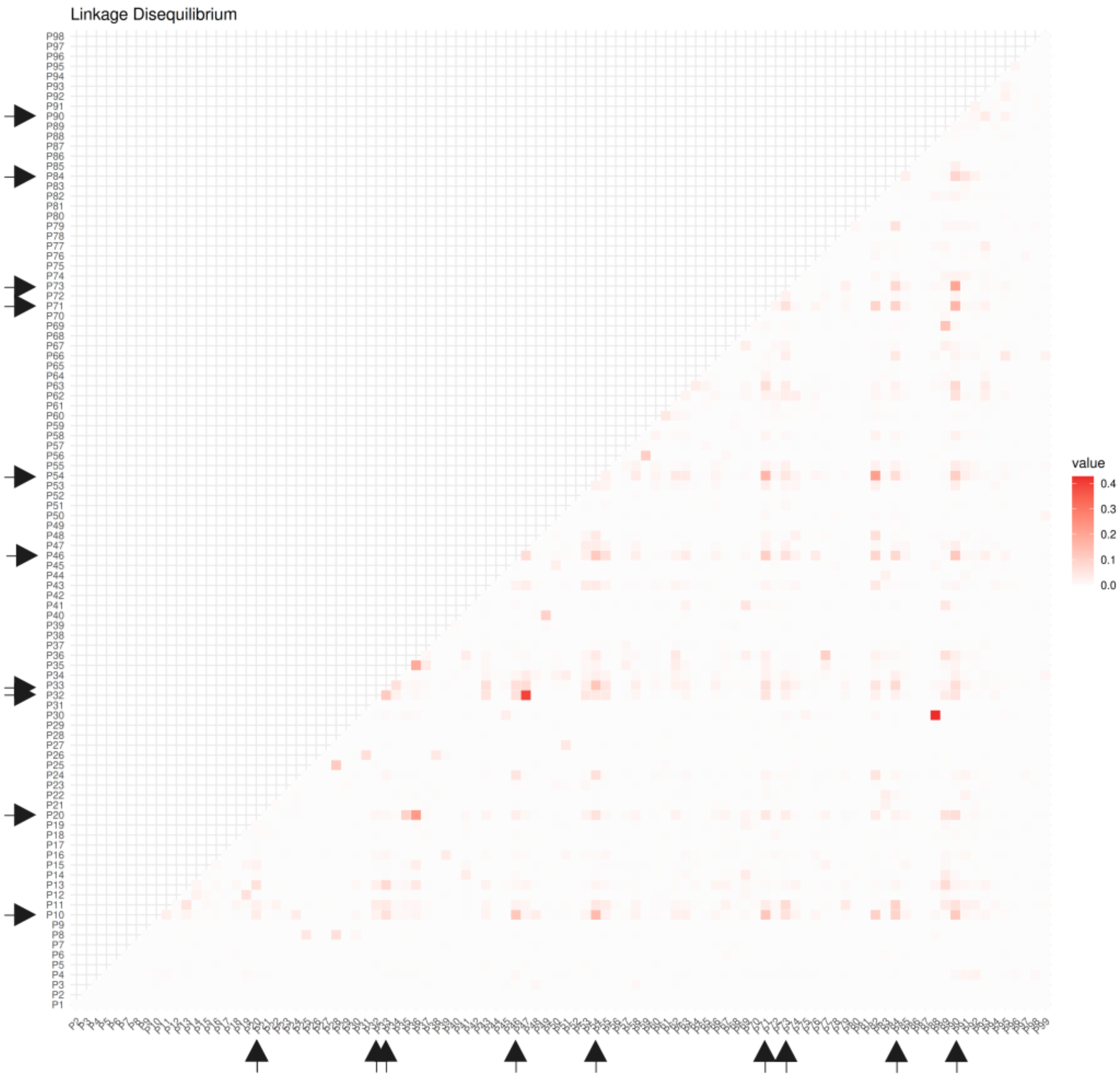


Supplementary Figure 1: Pairwise $r^2$ values between the 99 positions of protease. Stronger LD is represented in red. The top ten positions are highlighted with arrows.

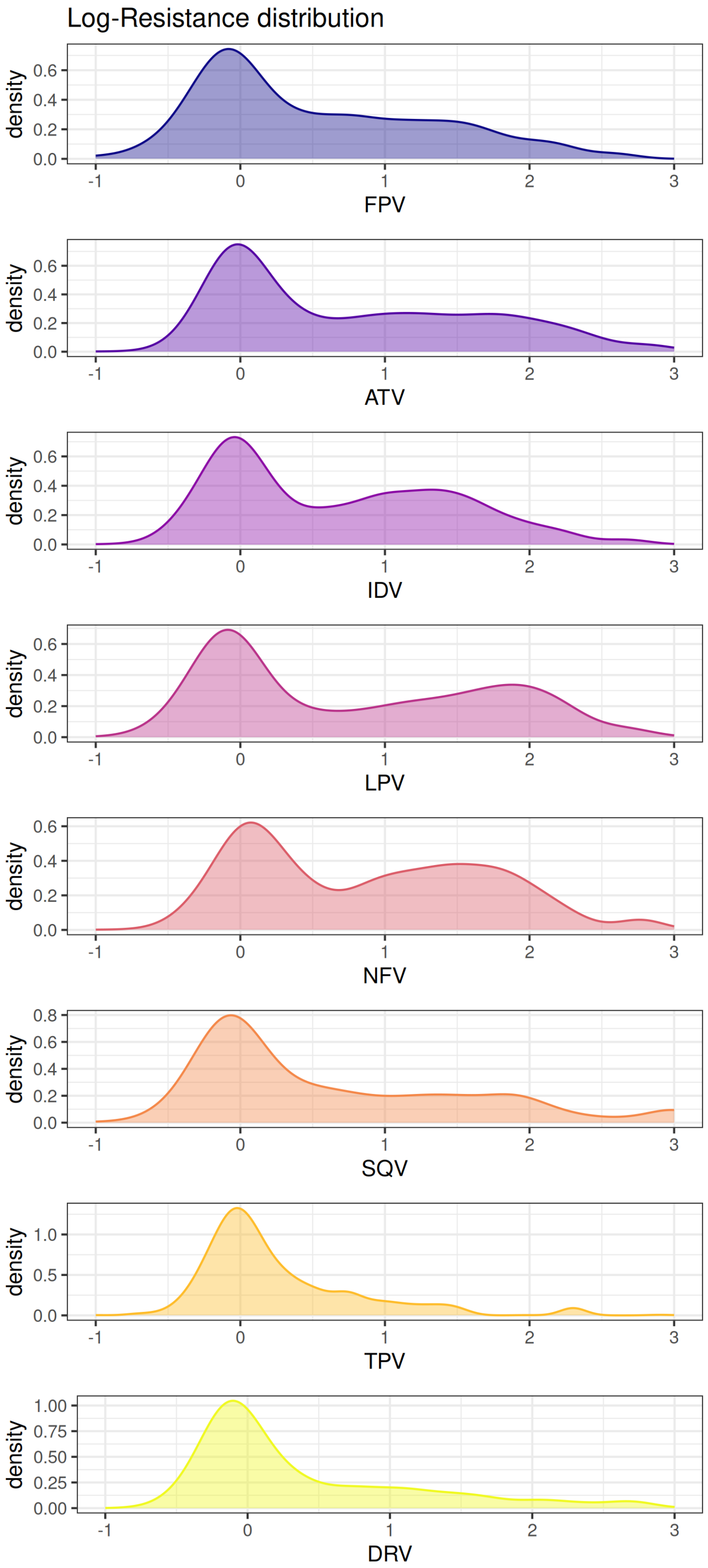


Supplementary Figure 2: log-IC50 density functions for the eight PIs.

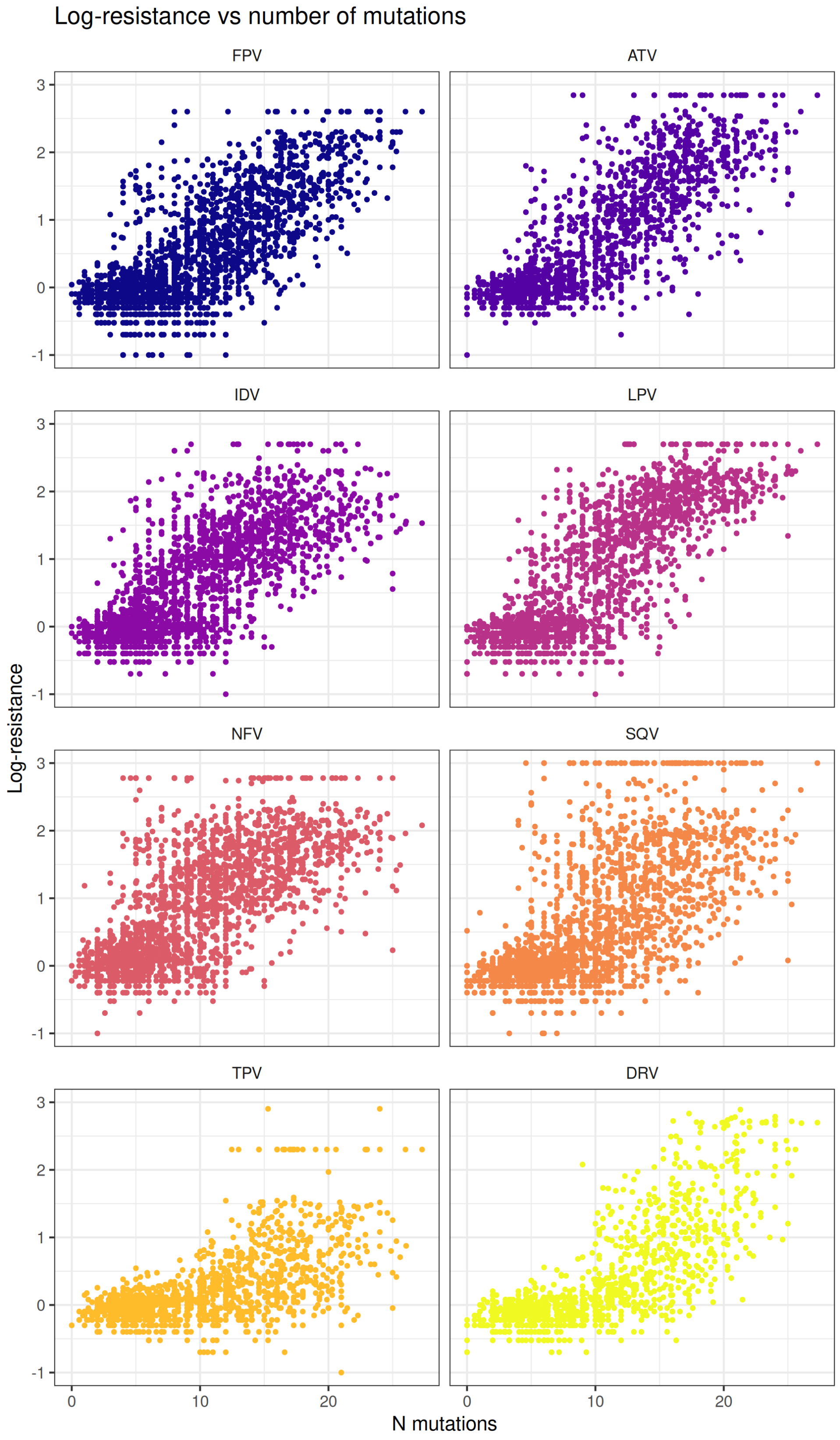


Supplementary Figure 3: Log-IC50 vs number of mutations (per isolate).

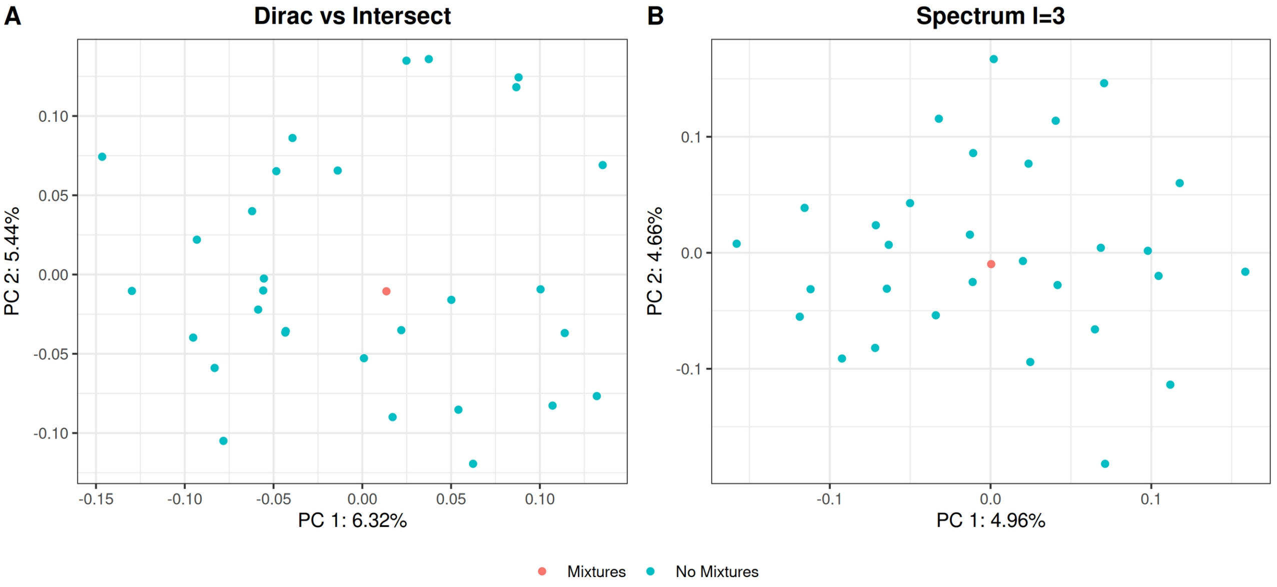


Supplementary Figure 4: Kendall's τ comparison among PC1 from the kernels computed from the original dataset (red) and the 30 resamples of the dataset without mixtures (turquoise blue). Panel A: Kernels for amino acid substitutions (Dirac vs Intersect). Panel B: Kernels for trimers (Spectrum vs mixSpectrum *k*=3).

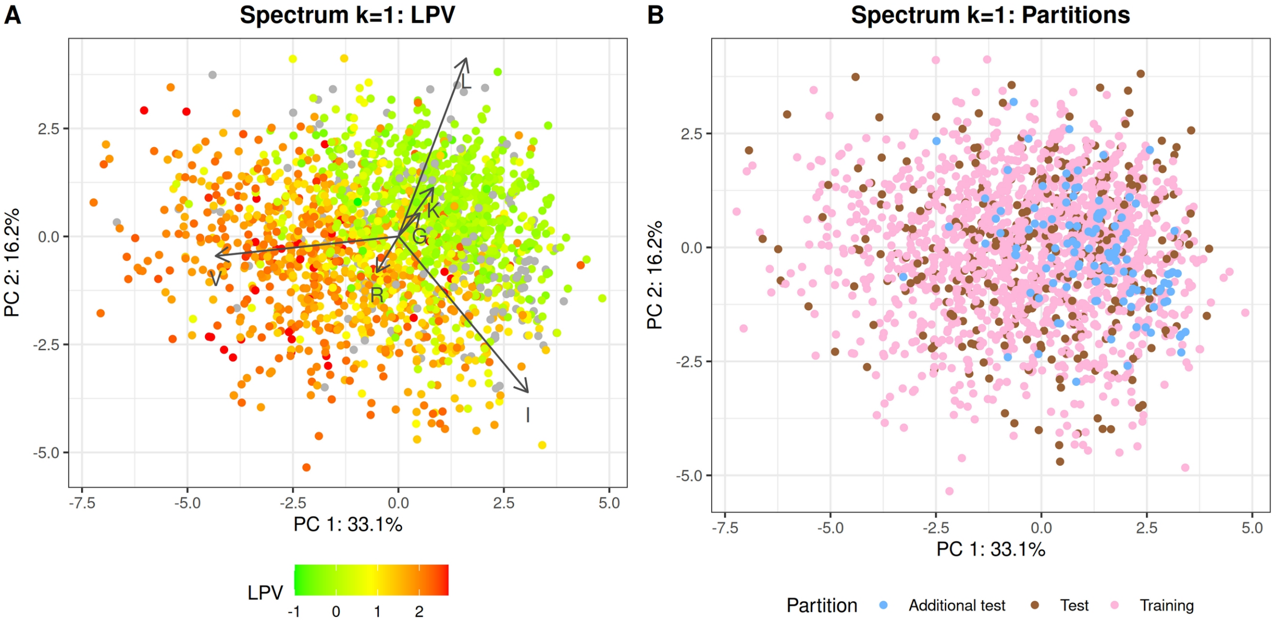


Supplementary Figure 5: Genotype and phenotype ordination according to Spectrum k=1 kPCA. Panel A: protease isolates colored according to their log-resistance value to LPV. Red means higher resistance, while green means higher susceptibility. Sequences with missing IC50 data for the LPV drug are in gray. Panel B: Training, test and additional test samples. Training samples are in pink and test samples in brown (both sets contain only HIV-1 subtype B isolates). Additional test samples (other subtypes) are in blue.

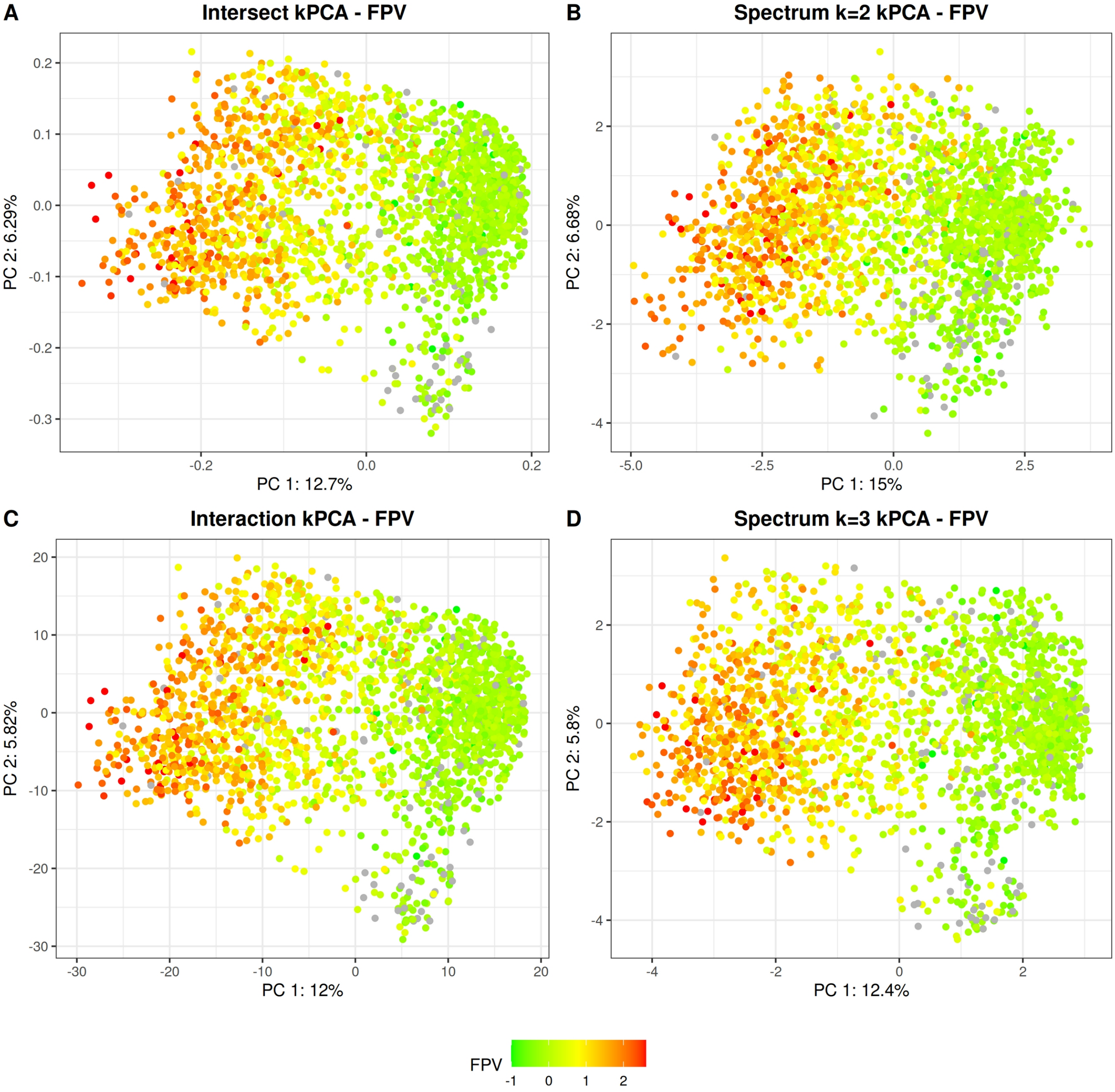


Supplementary Figure 6: Genotype vs phenotype ordination. Panel A: Intersect kPCA, panel B: Spectrum-2 kPCA, panel C: Interaction kPCA, panel D: Spectrum-3 kPCA. Each dot is a protease isolate, colored according to its log-resistance value to FPV. Red means higher resistance values, while green means higher susceptibility. Sequences with missing IC50 data for the drug are in gray.

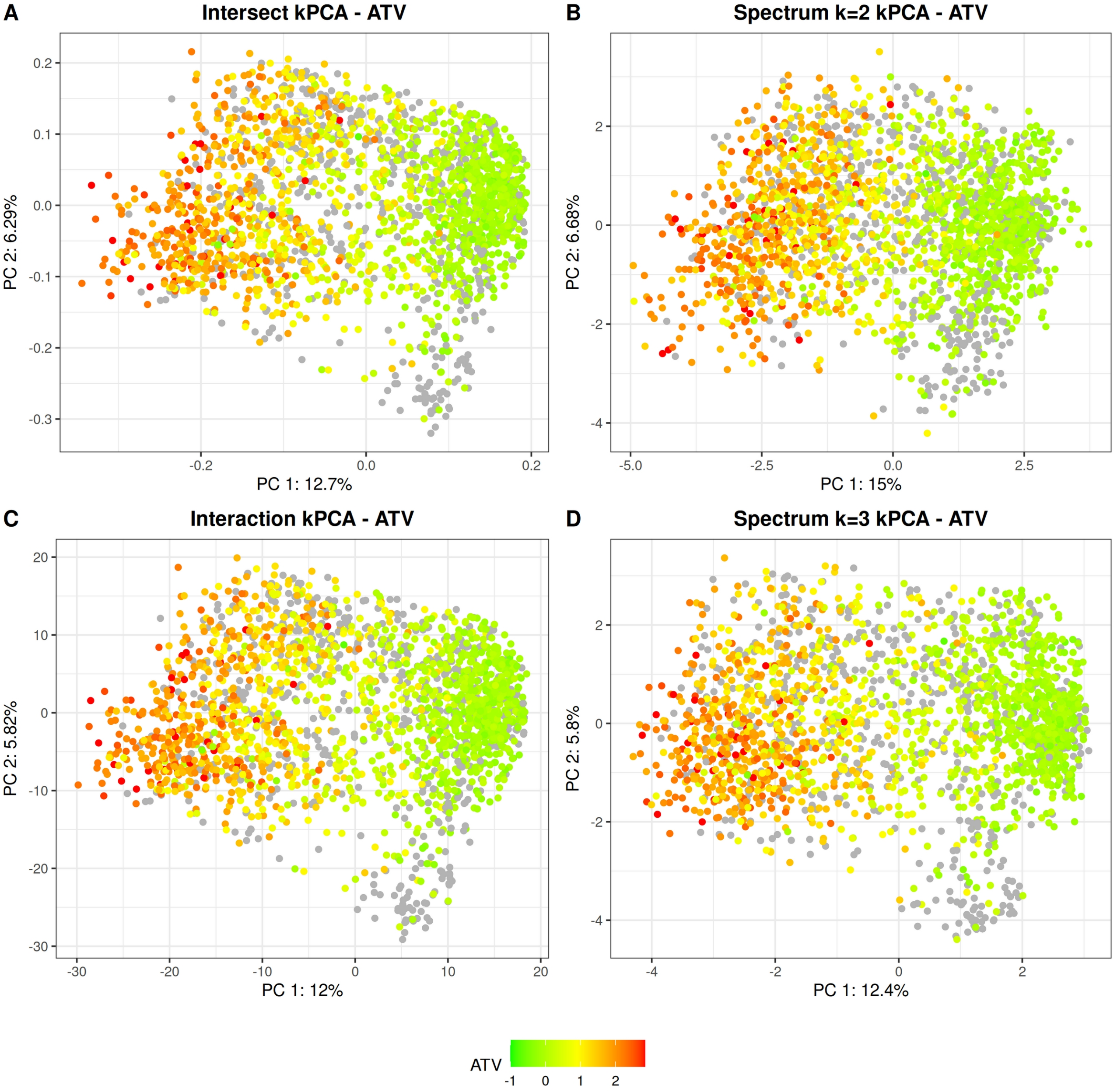


Supplementary Figure 7: Genotype vs phenotype ordination. Panel A: Intersect kPCA, panel B: Spectrum-2 kPCA, panel C: Interaction kPCA, panel D: Spectrum-3 kPCA. Each dot is a protease isolate, colored according to its log-resistance value to ATV. Red means higher resistance values, while green means higher susceptibility. Sequences with missing IC50 data for the drug are in gray.

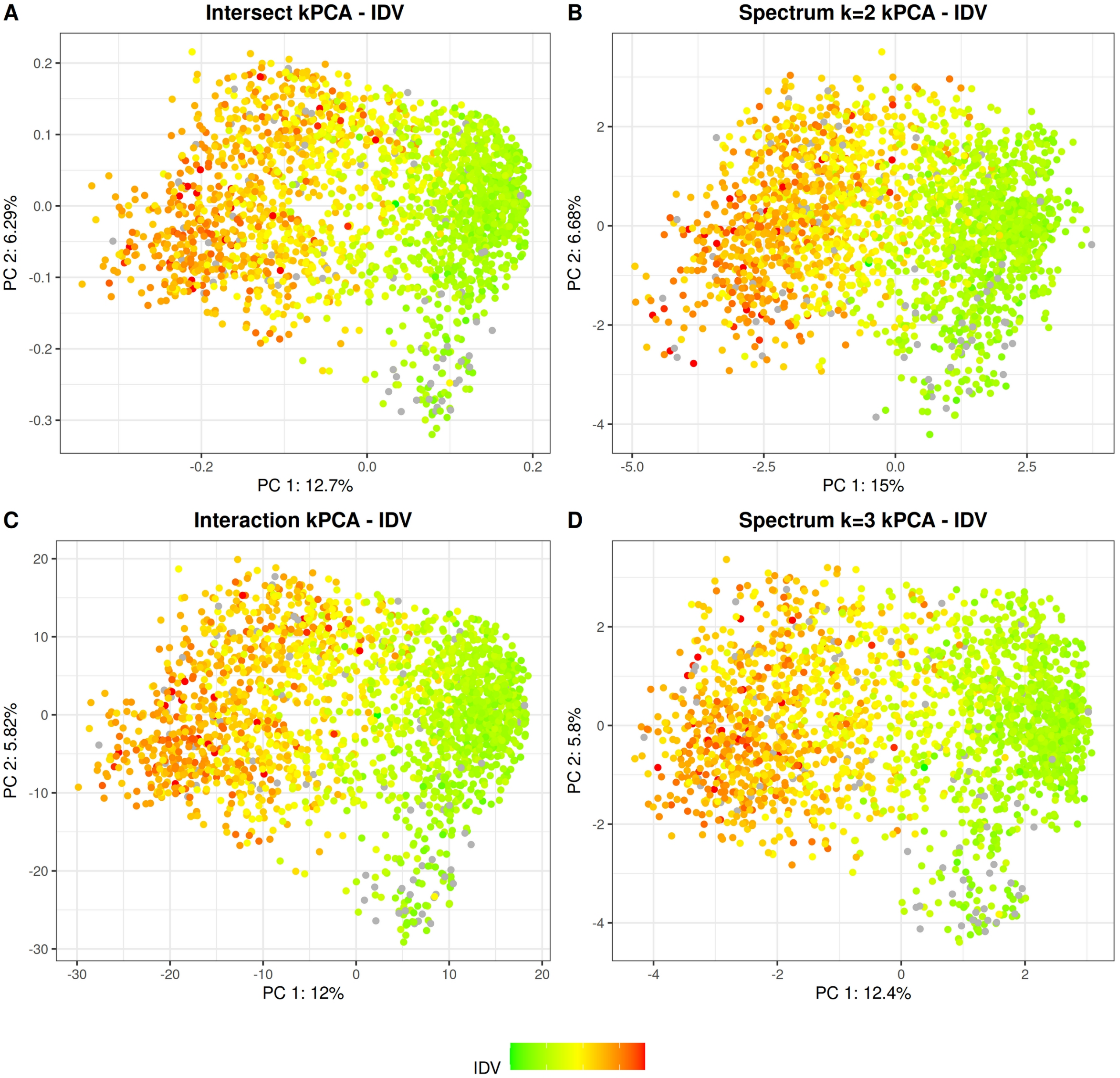


Supplementary Figure 8: Genotype vs phenotype ordination. Panel A: Intersect kPCA, panel B: Spectrum-2 kPCA, panel C: Interaction kPCA, panel D: Spectrum-3 kPCA. Each dot is a protease isolate, colored according to its log-resistance value to IDV. Red means higher resistance values, while green means higher susceptibility. Sequences with missing IC50 data for the drug are in gray.

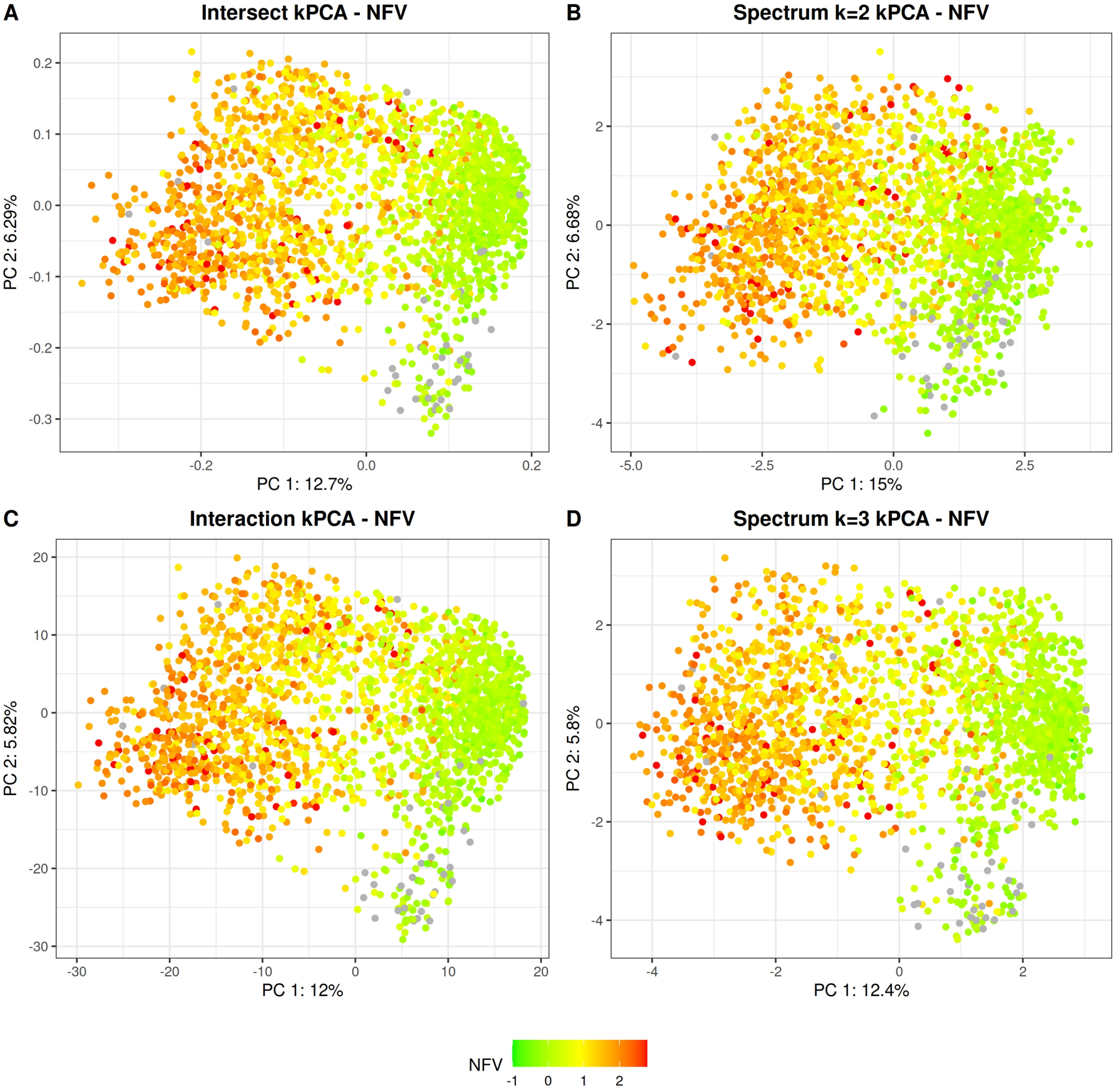


Supplementary Figure 9: Genotype vs phenotype ordination. Panel A: Intersect kPCA, panel B: Spectrum-2 kPCA, panel C: Interaction kPCA, panel D: Spectrum-3 kPCA. Each dot is a protease isolate, colored according to its log-resistance value to NFV. Red means higher resistance values, while green means higher susceptibility. Sequences with missing IC50 data for the drug are in gray.

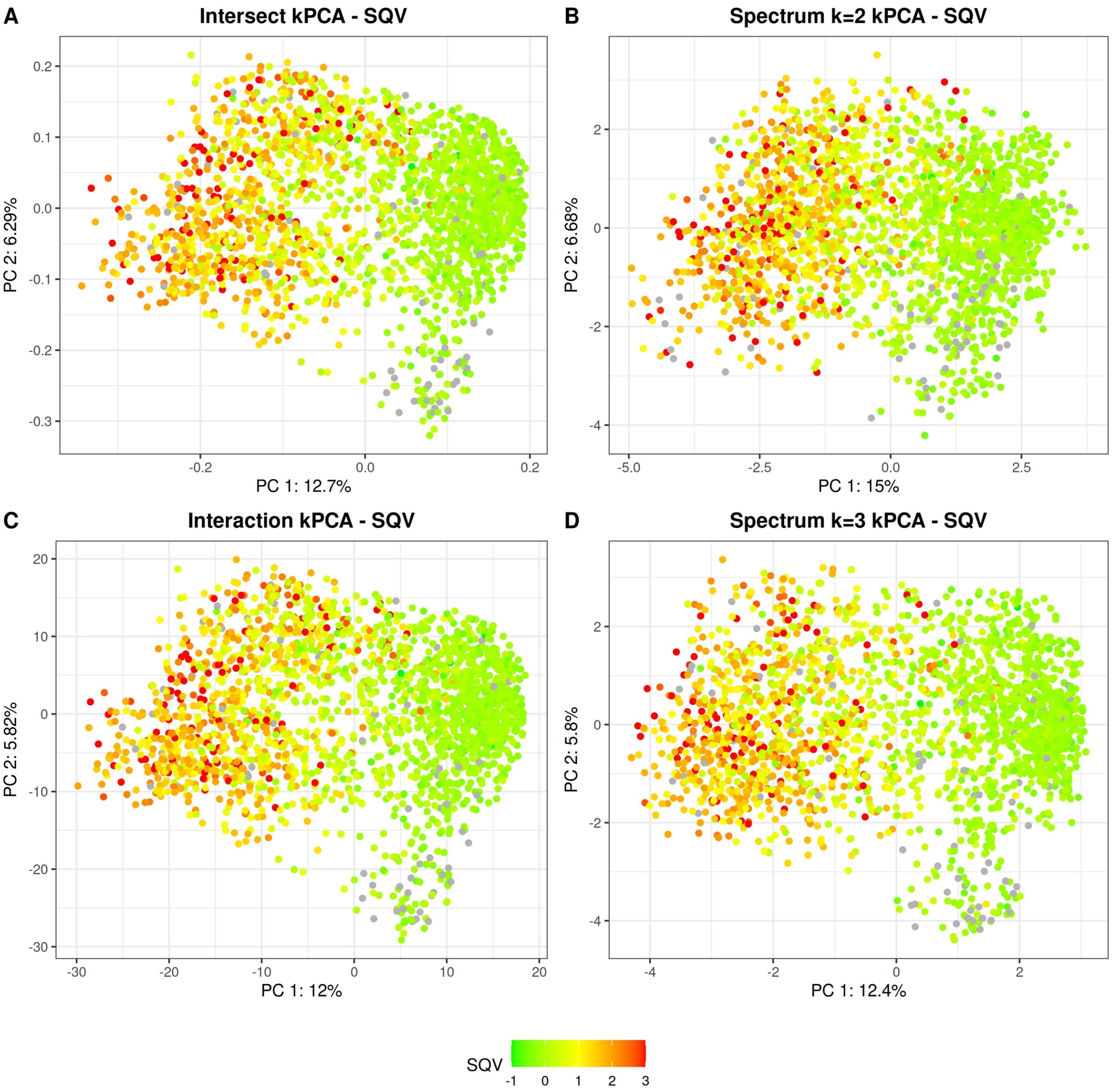


Supplementary Figure 10: Genotype vs phenotype ordination. Panel A: Intersect kPCA, panel B: Spectrum-2 kPCA, panel C: Interaction kPCA, panel D: Spectrum-3 kPCA. Each dot is a protease isolate, colored according to its log-resistance value to SQV. Red means higher resistance values, while green means higher susceptibility. Sequences with missing IC50 data for the drug are in gray.

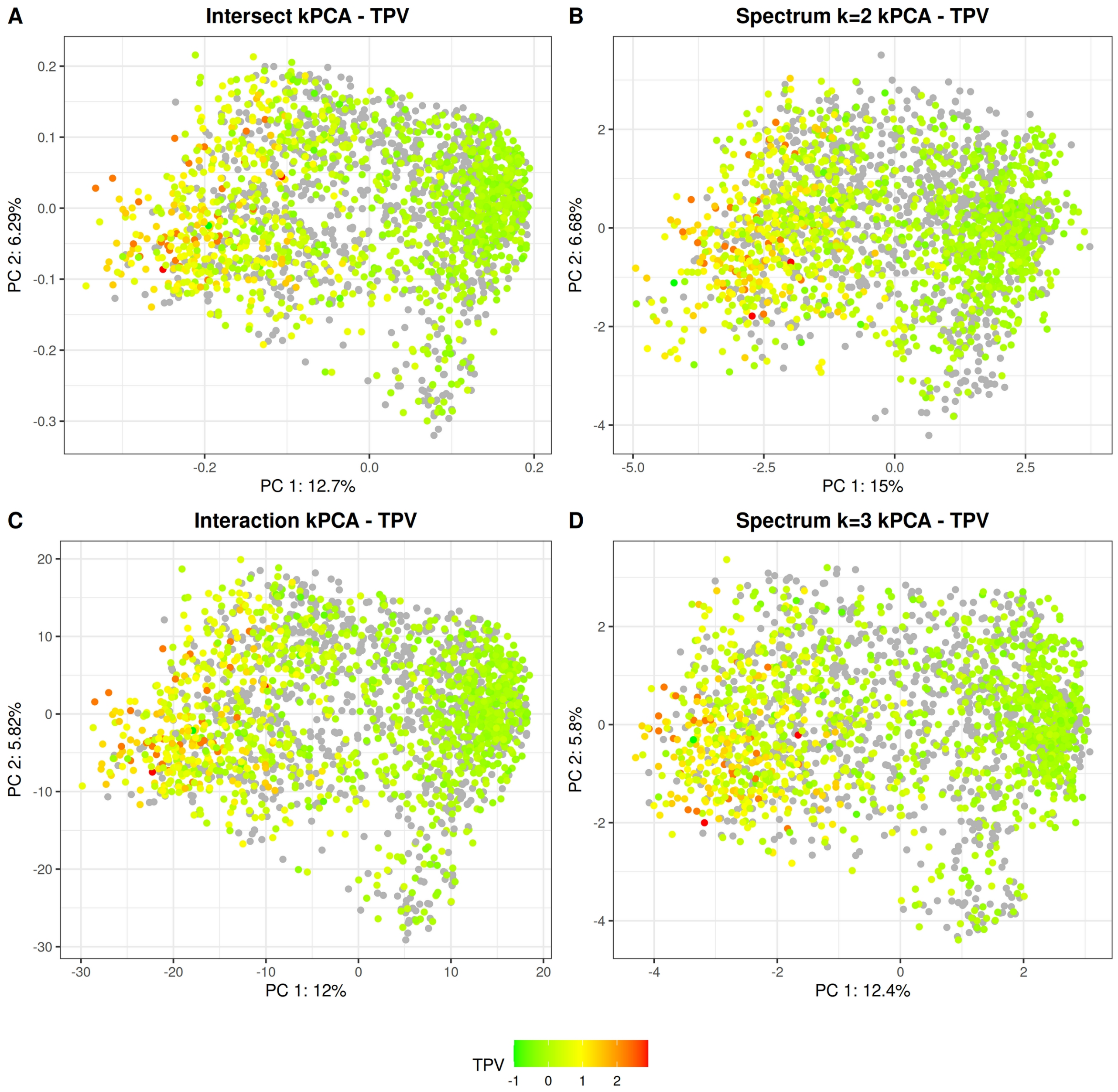


Supplementary Figure 11: Genotype vs phenotype ordination. Panel A: Intersect kPCA, panel B: Spectrum-2 kPCA, panel C: Interaction kPCA, panel D: Spectrum-3 kPCA. Each dot is a protease isolate, colored according to its log-resistance value to TPV. Red means higher resistance values, while green means higher susceptibility. Sequences with missing IC50 data for the drug are in gray.

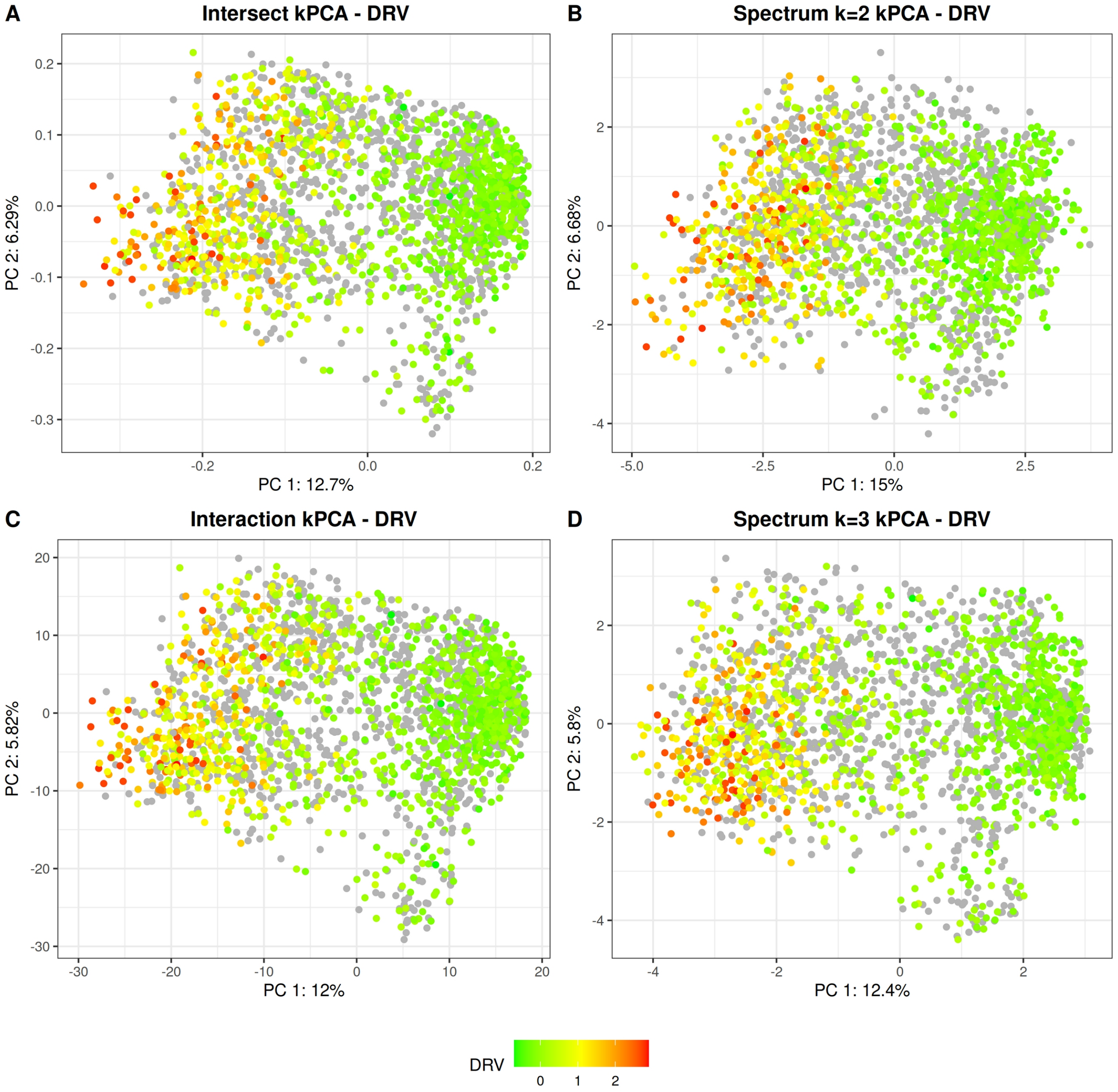


Supplementary Figure 12: Genotype vs phenotype ordination. Panel A: Intersect kPCA, panel B: Spectrum-2 kPCA, panel C: Interaction kPCA, panel D: Spectrum-3 kPCA. Each dot is a protease isolate, colored according to its log-resistance value to DRV. Red means higher resistance values, while green means higher susceptibility. Sequences with missing IC50 data for the drug are in gray.

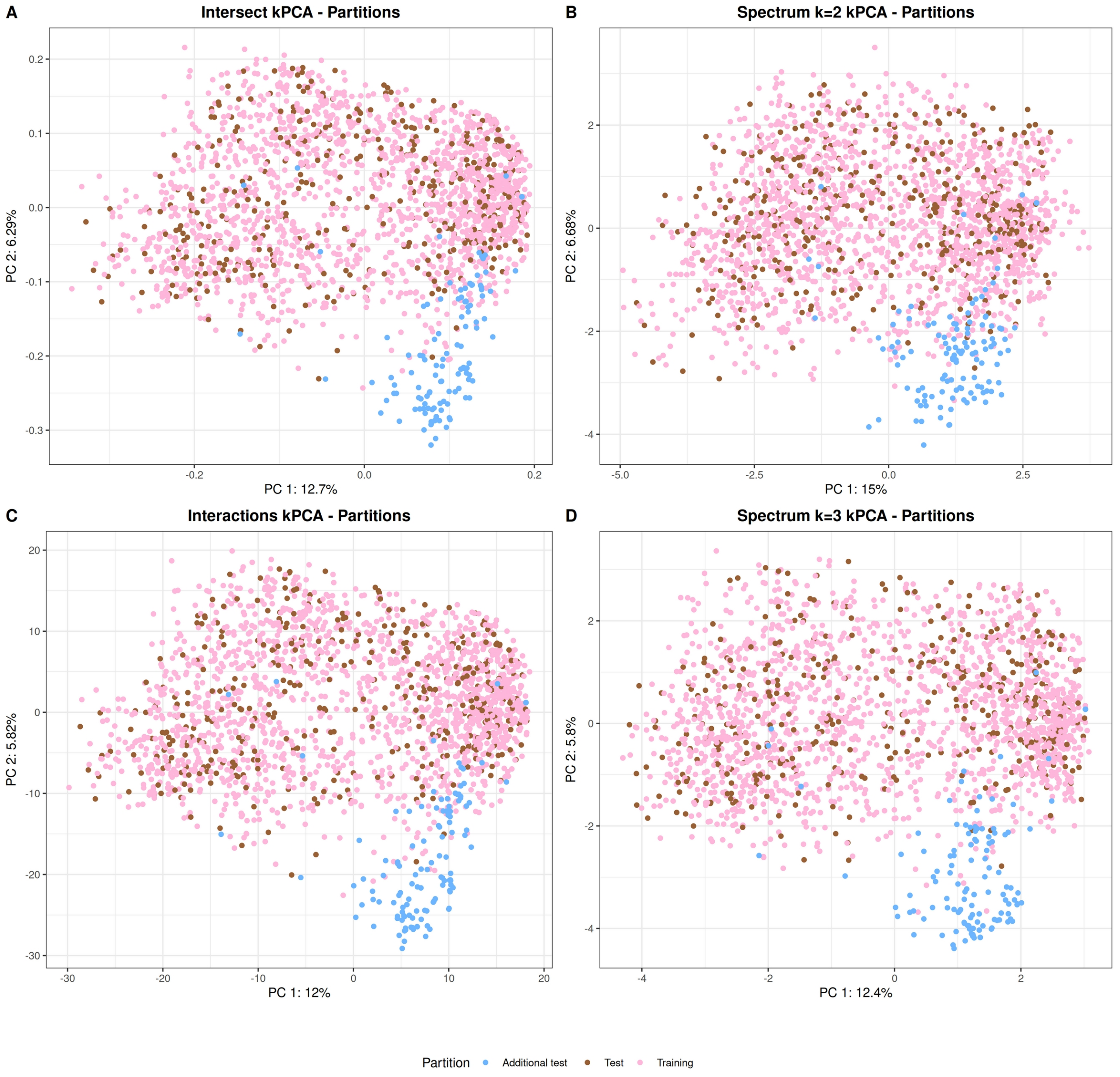


Supplementary Figure 13: Training, test and additional test samples. The kPCAs are identical to those of Figure 2. Panel A: Intersect kPCA, panel B: Spectrum-2 kPCA, panel C: Interaction kPCA, panel D: Spectrum-3 kPCA. Training samples are in pink and test samples in brown (both sets contain only HIV-1 subtype B isolates). Additional test samples (other subtypes) are in blue.

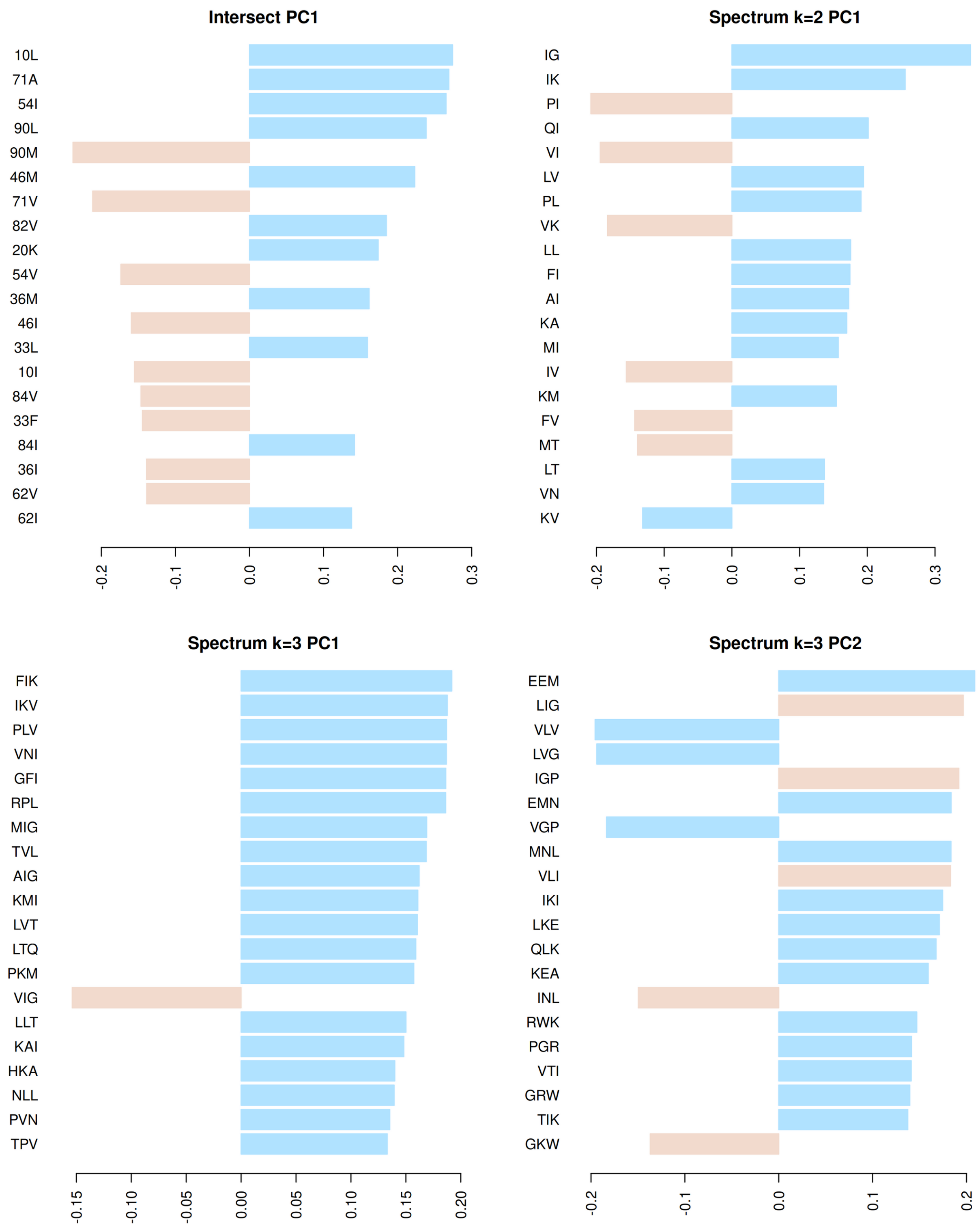


Supplementary Figure 14: Top-20 alleles with more contribution to the Intersect PC1 (panel A), Spectrum-2 PC1 (panel B), Spectrum-3 PC1 (panel C) and PC2 (panel D). The WT alleles are colored in blue, while the alternative alleles are in light brown.

Supplementary Table 1. SVM models' performance (computed via NMSE).

| | FPV | ATV | IDV | LPV | NFV | SQV | TPV | DRV |
|---|---|---|---|---|---|---|---|---|
| **Validation** | | | | | | | | |
| Spectrum-1 | 0.410 | 0.429 | 0.414 | 0.372 | 0.518 | 0.528 | 0.686 | 0.447 |
| Spectrum-2 | 0.136 | 0.170 | 0.152 | 0.111 | 0.176 | 0.207 | 0.347 | 0.142 |
| Spectrum-3 | 0.128 | 0.137 | 0.130 | 0.093 | 0.162 | 0.161 | 0.298 | 0.126 |
| Intersect | 0.119 | 0.128 | 0.124 | 0.082 | 0.142 | 0.148 | 0.288 | 0.112 |
| Interaction | 0.113 | 0.123 | 0.109 | 0.079 | 0.125 | 0.140 | 0.276 | 0.106 |
| **Test** | | | | | | | | |
| Spectrum1 | 0.473 | 0.367 | 0.439 | 0.349 | 0.509 | 0.542 | 0.670 | 0.501 |
| Spectrum2 | 0.140 | 0.167 | 0.173 | 0.120 | 0.186 | 0.222 | 0.322 | 0.131 |
| Spectrum3 | 0.129 | 0.154 | 0.131 | 0.105 | 0.198 | 0.150 | 0.326 | 0.119 |
| Intersect | 0.111 | 0.134 | 0.132 | 0.093 | 0.156 | 0.141 | 0.263 | 0.119 |
| Interaction | 0.113 | 0.128 | 0.120 | 0.093 | 0.137 | 0.127 | 0.244 | 0.119 |
| N test | 391 | 308 | 394 | 350 | 402 | 397 | 237 | 215 |
| **Additional test** | | | | | | | | |
| Spectrum-1 | 0.765 | 0.555 | 1.869 | 1.079 | 1.581 | 1.970 | 0.756 | 0.475 |
| Spectrum-2 | 0.545 | 0.286 | 0.669 | 0.315 | 0.858 | 1.054 | 0.850 | 0.217 |
| Spectrum-3 | 0.468 | 0.262 | 0.836 | 0.386 | 1.245 | 0.799 | 1.189 | 0.193 |
| Intersect | 0.535 | 0.241 | 0.573 | 0.351 | 0.723 | 0.618 | 1.017 | 0.221 |
| Interaction | 0.447 | 0.287 | 0.473 | 0.325 | 0.680 | 0.698 | 1.302 | 0.263 |
| N additional test | 90 | 36 | 90 | 79 | 91 | 95 | 49 | 53 |

Sets sizes are included as an additional information. Validation values correspond to the average of the 5 outer loops in 5x2 nested cross-validation. The test and additional test errors presented here are the "raw values" without bootstrapping.

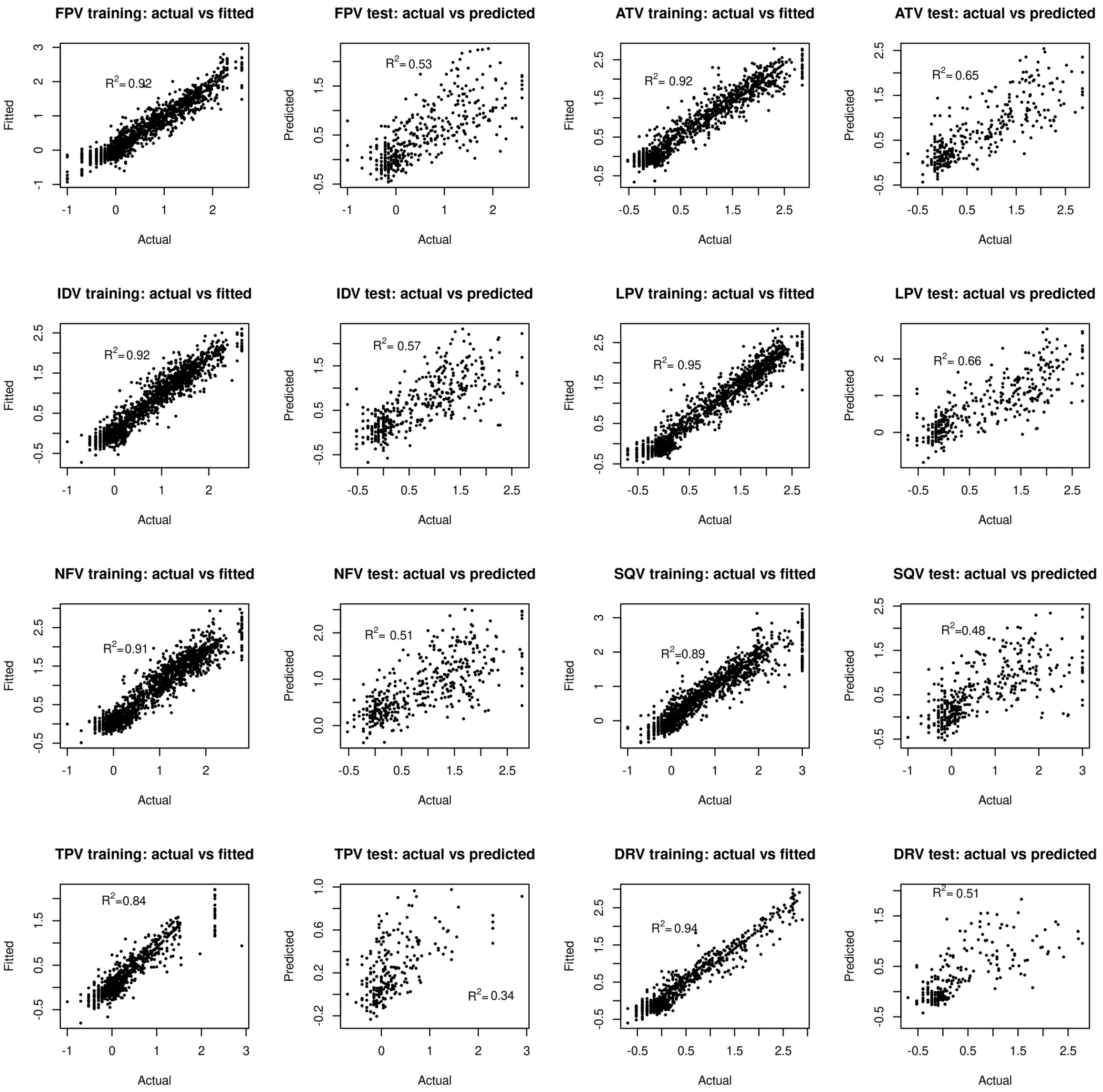


Supplementary Figure 15: mixSpectrum k = 1 kernel SVM model. Actual values vs fitted (in training) or predicted (test).

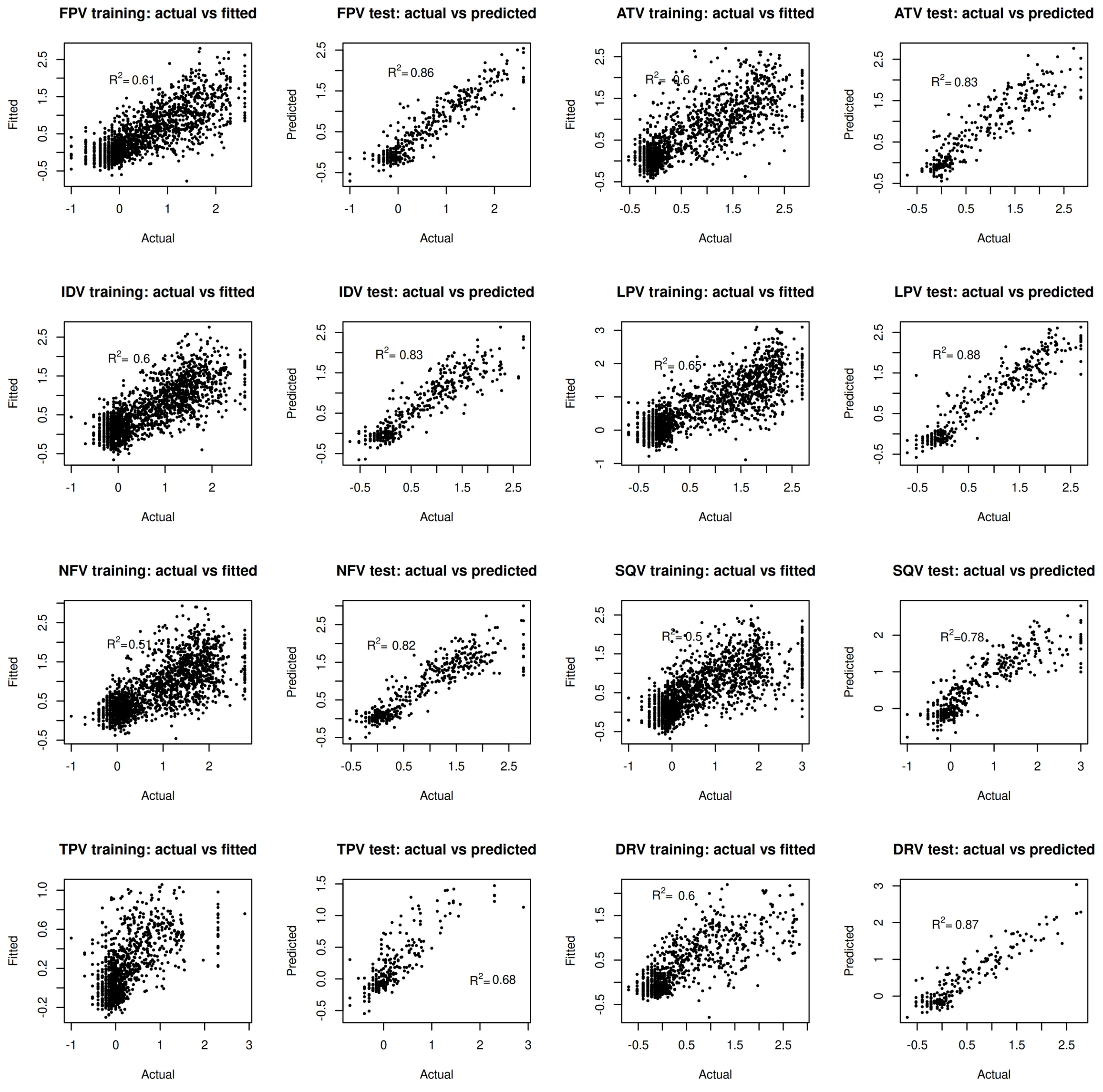


Supplementary Figure 16: mixSpectrum k = 2 kernel SVM model. Actual values vs fitted (in training) or predicted (test).

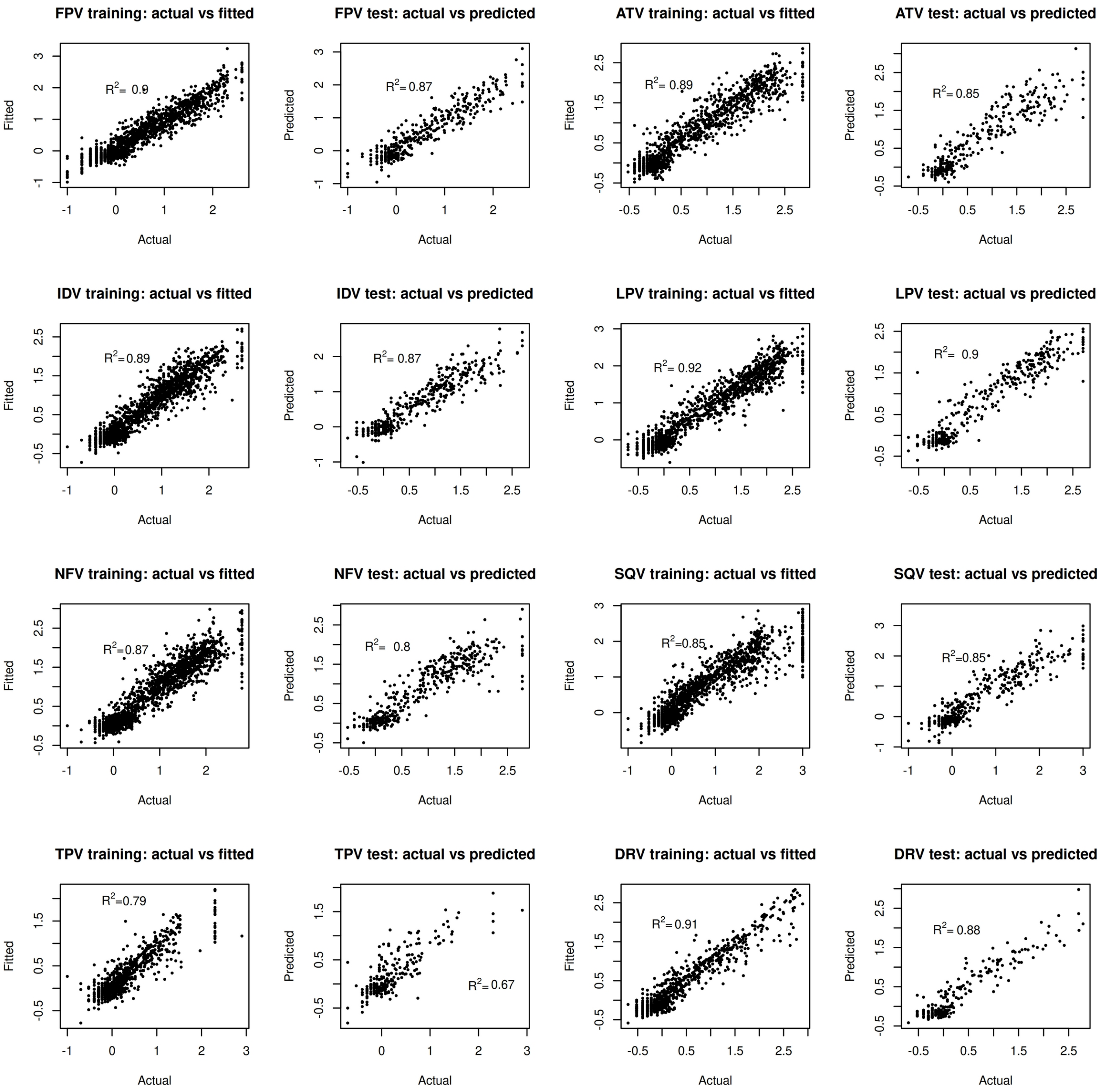


Supplementary Figure 17: mixSpectrum k = 3 kernel SVM model. Actual values vs fitted (in training) or predicted (test).

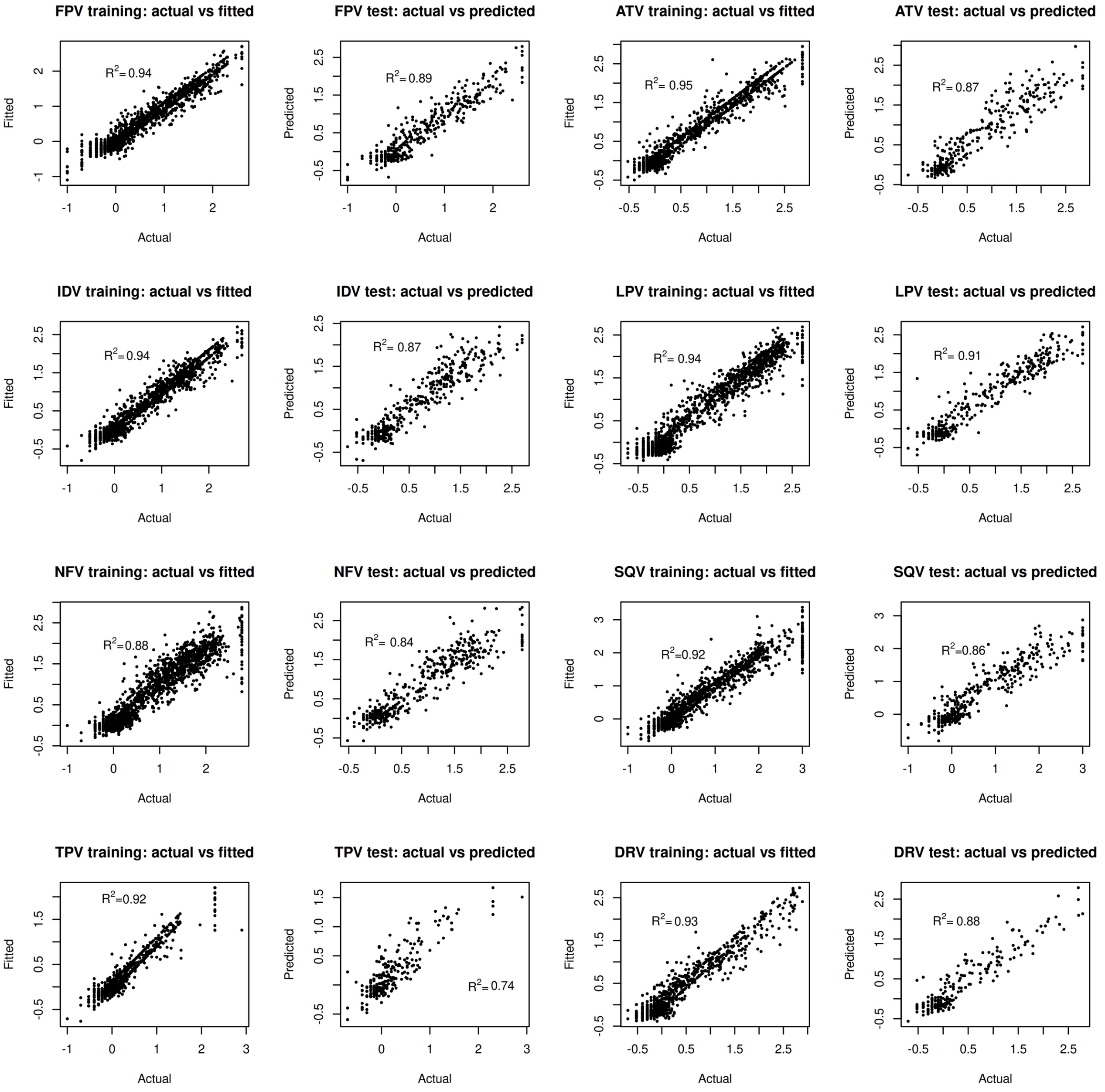


Supplementary Figure 18: Intersect kernel SVM model. Actual values vs fitted (in training) or predicted (test).

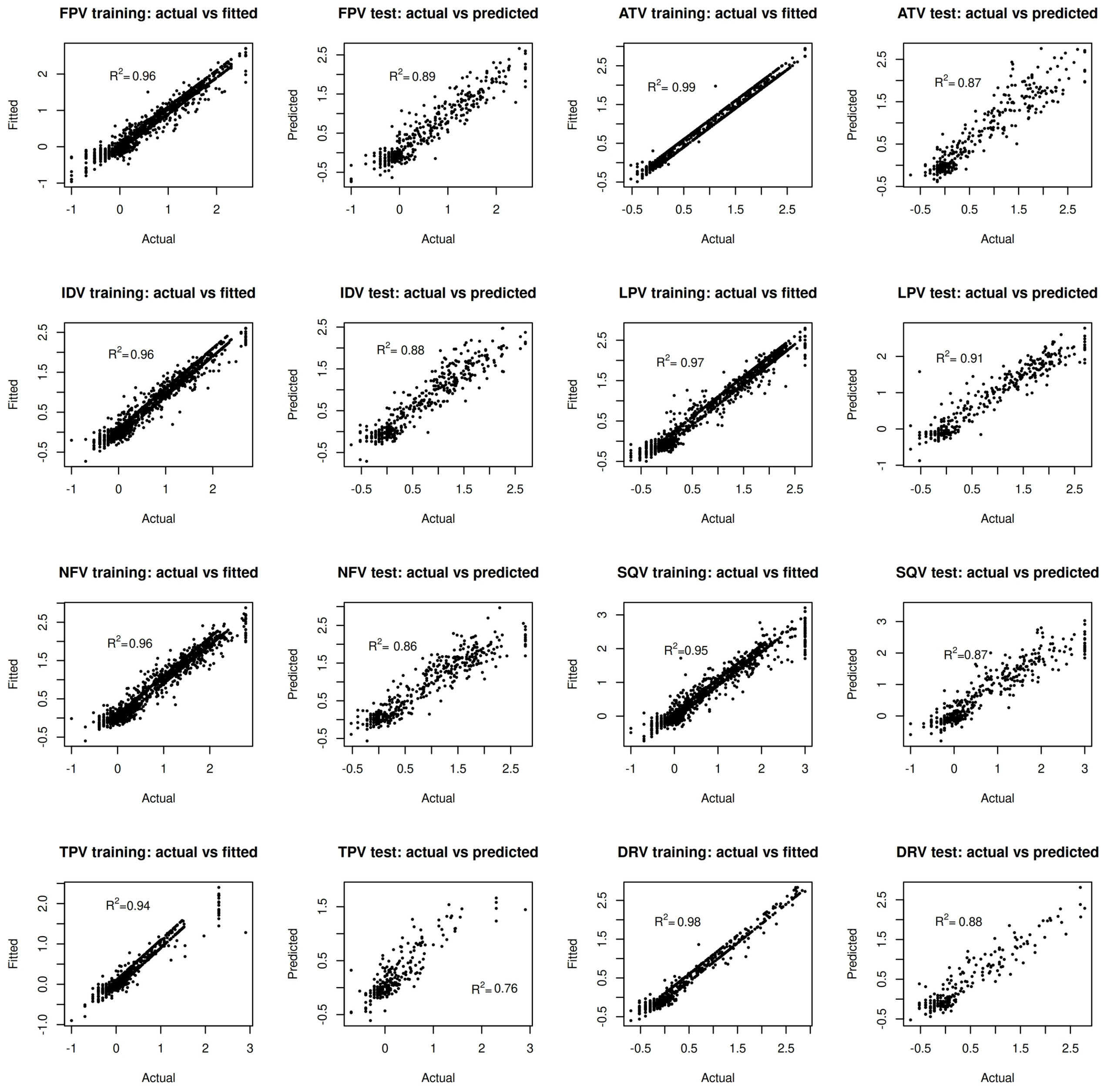


Supplementary Figure 19: Interaction kernel SVM model. Actual values vs fitted (in training) or predicted (test).

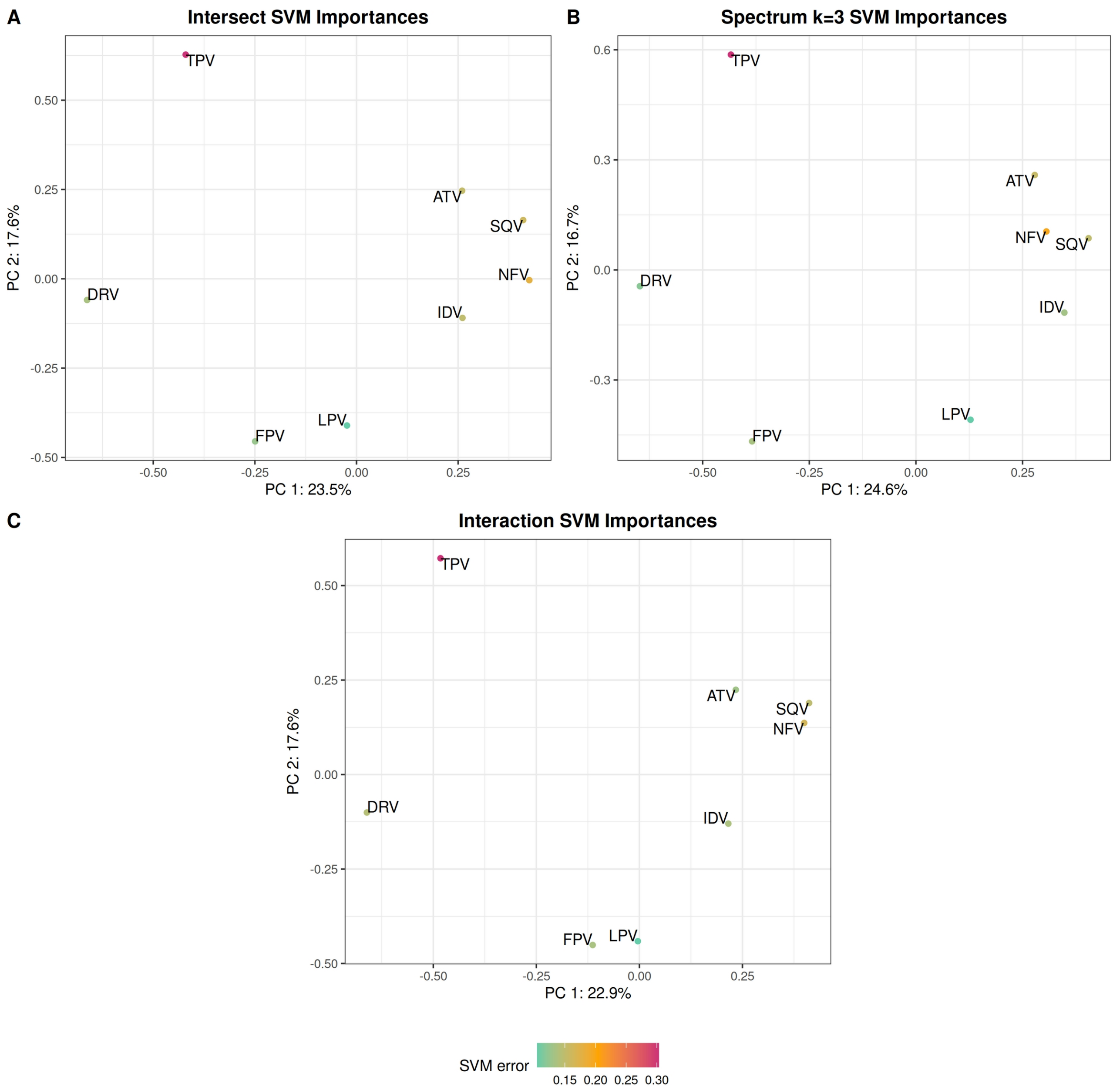


Supplementary Figure 20: Kendall's τ PCA comparing the eight SVM models (one for each drug) according to their feature rankings (which are single amino acid residues in Intersect kernel PCA, trimers in Spectrum-3, and pairwise interactions in Interaction). Color represents the NMSE of each model (see Supplementary Table 1).
